# LDB1 establishes multi-enhancer networks to regulate gene expression

**DOI:** 10.1101/2024.08.23.609430

**Authors:** Nicholas G. Aboreden, Jessica C. Lam, Viraat Y. Goel, Siqing Wang, Xiaokang Wang, Susannah C. Midla, Alma Quijano, Cheryl A. Keller, Belinda M. Giardine, Ross C. Hardison, Haoyue Zhang, Anders S. Hansen, Gerd A. Blobel

**Affiliations:** Perelman School of Medicine, University of Pennsylvania, Philadelphia, PA, USA; Division of Hematology, The Children’s Hospital of Philadelphia, Philadelphia, PA, USA; Department of Biological Engineering, Massachusetts Institute of Technology, Cambridge, MA, USA; Gene Regulation Observatory, Broad Institute of MIT and Harvard, Cambridge, MA, USA; Koch Institute for Integrative Cancer Research, Cambridge, MA, USA; Department of Biochemistry and Molecular Biology, Pennsylvania State University, University Park, PA, USA; Institute of Molecular Physiology, Shenzhen Bay Laboratory, Shenzhen, Guangdong, China

## Abstract

How specific enhancer-promoter pairing is established is still mostly unclear. Besides the CTCF/cohesin machinery, only a few nuclear factors have been studied for a direct role in physically connecting regulatory elements. Here, we show via acute degradation experiments that LDB1 directly and broadly promotes enhancer-promoter loops. Most LDB1-mediated contacts, even those spanning hundreds of kb, can form in the absence of CTCF, cohesin, or YY1 as determined via the use of multiple degron systems. Moreover, an engineered LDB1-driven chromatin loop is cohesin independent. Cohesin-driven loop extrusion does not stall at LDB1 occupied sites but may aid the formation of a subset of LDB1 anchored loops. Leveraging the dynamic reorganization of nuclear architecture during the transition from mitosis to G1-phase, we establish a relationship between LDB1-dependent interactions in the context of TAD organization and gene activation. Lastly, Tri-C and Region Capture Micro-C reveal that LDB1 organizes multi-enhancer networks to activate transcription. This establishes LDB1 as a direct driver of regulatory network inter-connectivity.

## INTRODUCTION

Cell-type-specific gene expression signatures rely on the action of enhancers which can act over large genomic distances and do not always regulate the nearest gene^1,2^. Long range action by most enhancers is achieved by physical proximity with promoters, highlighting the intricate connection between genome architecture and transcription regulation^3–7^.

Enhancer-promoter (E-P) connectivity is influenced by sub-megabase scale topologically associating domains^8–12^ (TADs) which favor regulatory contacts within their boundaries and disfavor contacts across them. TAD boundaries are frequently co-occupied by CTCF and cohesin. The ring-like cohesin complex is believed to extrude the chromatid until it is stalled by convergently oriented CTCF sites, resulting in transient looped contacts, also referred to as structural loops^13–16^. Hence, TADs encompassed by CTCF/cohesin loops are also called loop domains.

The organization of CTCF/cohesin loops can impact E-P connectivity in multiple ways. CTCF is known to function as an enhancer-blocking insulator when positioned between an enhancer and promoter, and its insulation function is linked to its ability to form chromatin loops^17–25^. On the other hand, E-P contacts can be supported by structural loops, especially when the E-P loop anchors are closely flanked by the structural loop anchors^26,27^. In addition, chromatin extrusion intermediates may facilitate the probability of E-P encounters that are subsequently maintained by promoter- and enhancer-bound transcription (co-)factors. This seems to be especially the case for long-range E-P contacts that may be more dependent on cohesin than short range ones^28,29^.

Depletion of CTCF or cohesin abrogates most loop domains, yet the effects on gene expression are surprisingly mild^27,30–35^ implying that most regulatory connectivity remains intact in their absence. Furthermore, E-P contacts can be established prior to structural loop formation during the mitosis-to-G1 phase transition, an interval during which randomly looped chromatin is re-organized into interphase-like state in newborn daughter nuclei^36^. Many such E-P contacts can even be rebuilt in the absence of CTCF^26^. More recently, the development of Region Capture Micro-C revealed that short-range and highly nested contacts between *Cis*-regulatory elements (CREs) remain intact following cohesin depletion^37^.

In concert, these studies suggest that CTCF and cohesin may influence E-P connectivity in a context-dependent manner but that their requirement is not absolute. How CTCF/cohesin independent long range contacts are formed, and which factors convey specificity remain critical questions in the field.

The advent of acute degradation technologies has enabled the interrogation of direct or proximal roles of individual proteins in mediating CRE contacts^38^, including those of the CTCF and cohesin machinery. While numerous factors have been implicated in CRE connectivity^39–53^, few have been studied using an acute depletion strategy, leaving open the possibility that observations may be due to cell fate changes or other secondary effects. For example, in the case of YY1, a transcription factor with architectural roles, long term (24 hr) depletion had a much more profound effect on CRE connectivity when compared to acute (3 hr) depletion^39,54,55^. More generally, the identification and mechanistic studies of transcription factors that directly control long-range CRE interactions as determined by short-term depletion, and how they may be influenced by the process of loop extrusion has lagged behind studies on loops formed by the CTCF/cohesin machinery.

Mounting evidence supports a role for the transcription co-factor LDB1 as an architectural protein. LDB1 does not bind DNA directly but is recruited to CREs via tissue-specific DNA binding proteins such as the erythroid transcription factors GATA1 and TAL1^56–65^. Loss- and gain-of-function experiments at the β-globin locus implicate LDB1 as a mediator of enhancer-promoter proximity^40,66–68^. At this locus, LDB1 may establish a homotypic looping interaction by occupying both enhancer and β-globin promoter elements. However, at different loci heterotypic interactions (in which LDB1 occupies only one interacting element) have also been proposed^69^. LDB1-dependent contacts at the β-globin locus can be established in the absence of focal cohesin accumulation, suggesting that LDB1 does not function as a cohesin stalling factor at this locus, but neither rules out such a function at other loci nor does address a possible role of cohesin extrusion intermediates as facilitators of LDB1-mediated contacts^70^. Additional studies indicate LDB1’s involvement in enhancer/promoter connectivity in diverse cell types^71–73^. For example, in post-mitotic neurons, LDB1 is required for the maintenance of both intra- and inter-chromosomal contacts^74^. While these elegant studies strongly suggest a role for LDB1 in regulatory connectivity, none of them are immune to the caveats intrinsic to prolonged perturbations such as potentially confounding secondary effects. Moreover, none of these prior studies explored any mutual influence of LDB1-driven and CTCF, cohesin or YY1-driven forces that shape the mammalian genome.

Here, we employed an acute degradation system and exploited cell cycle dynamics in combination with Micro-C^75–77^, Region Capture Micro-C^37^, and Tri-C^78^ to comprehensively study LDB1’s direct role in chromatin architecture and transcription. We find that LDB1 is required to maintain wide-spread chromatin contacts between CREs. LDB1 organizes complex, multi-enhancer networks that can involve extremely short-range contacts. Importantly, there is minimal overlap between LDB1-dependent loop anchors and CTCF/cohesin genome wide, arguing against LDB1 functioning as a loop extrusion barrier. By integrating CTCF, cohesin and YY1 degradation experiments, we found that the majority of LDB1-driven contacts do not rely on CTCF or cohesin. However, the cohesin mediated extrusion process may assist in the formation of a subset of LDB1 dependent loops. Our findings highlight LDB1 as an important mechanistic link between chromatin architecture and transcriptional regulation. We suggest that enhancer/promoter communication is simultaneously achieved through specific and generic forces; the former represented by LDB1 mediated contacts, and the latter by general architectural factors that may promote or constrain them.

## RESULTS

### LDB1 mediates chromatin contacts between cis-regulatory elements (CREs)

To test the direct role of LDB1 in chromatin architecture genome-wide, we tagged endogenous LDB1 homozygously with a minimal auxin-inducible degron (mAID) and mCherry via CRISPR-mediated gene editing in the G1E-ER4^79^ erythroblast cell line (Figure S1A). Upon 4 hours of auxin treatment, LDB1 was virtually completely degraded as measured by Western blot in cell lysates and by flow cytometry (Figure S1B, C). Since protein removal from chromatin can be uneven or delayed^80^ we carried out anti-LDB1 ChIP-seq which showed that the vast majority of LDB1 peaks were lost at this time point (Figure S1D). We next examined whether the mAID-mCherry tag interferes with LDB1 function by measuring the fusion protein’s ability to induce the expression of two erythroid LDB1 target genes: β-globin and Gypa^57^ in G1E-ER4 cells. G1E-ER4 cells are derived from the GATA1 null erythroblast cell line G1E and express GATA1 fused to the ligand binding domain of the estrogen receptor (GATA1-ER). Upon estradiol treatment, GATA1-ER activates numerous erythroid genes including β-globin in an LDB1-dependent manner^79^. LDB1-AID-mCherry clonal lines exhibited comparable levels of β-globin and Gypa activation to parental cells. Importantly, auxin treatment abrogated β-globin and Gypa activation in two independent clonal lines (Figure S1E). To further validate that the mAID-mCherry tag does not interfere with LDB1 function, we performed RNA-seq in parental G1E-ER4 cells and the same two LDB1-AID-mCherry clonal lines. We found a high concordance amongst the transcriptomes of each clonal line and the parental line demonstrating that the mAID-mCherry tag does not significantly alter gene expression profiles (Figure S1F-I). While both clones displayed comparable gene expression profiles relative to the parental line, clone 2 showed the highest consistency. Therefore, we selected clone 2 for subsequent experiments.

To measure the immediate consequences of LDB1 depletion on chromatin architecture, we performed Micro-C with or without 4 hours of auxin treatment. 9 biological replicates were pooled to attain ∼1.085 and ∼1.068 billion valid cis contacts for untreated and auxin-treated samples respectively (Supplementary table 1 and Figure S1J). The effects of LDB1 depletion on chromatin architecture were largely restricted to chromatin loops while compartments and TADs were minimally impacted upon LDB1 removal (Figure S1K-L). We identified a total of 20,926 chromatin loops as a unified list from both untreated and auxin-treated samples using the cooltools.dots function. We quantified loop strength for each loop per treatment condition by measuring the observed contact frequency within the peak pixel divided by a locally adjusted expected value. We assigned a log2FC value comparing LDB1 replete and depleted conditions for each loop and used a log2FC cutoff of −/+ 0.5 to define weakened and strengthened loops (e.g. weakened loops defined as at least a ∼30% reduction in loop strength).

To characterize the looping interactions controlled by LDB1, we classified loops based on whether loop anchors were located at CREs based on our prior annotations^36^. Loop anchors were defined as 10kb regions centered on the respective midpoints of the pixel encompassing all bin-bin pairs in the loop center. We also classified loops based on the presence of LDB1, CTCF and cohesin (RAD21) at one or both loop anchors. To that end, we generated ChIP-seq data sets for LDB1, CTCF, and RAD21 in our degron cell line (Figure 1M) and integrated these data with our chromatin loop calls.

**Figure 1.**
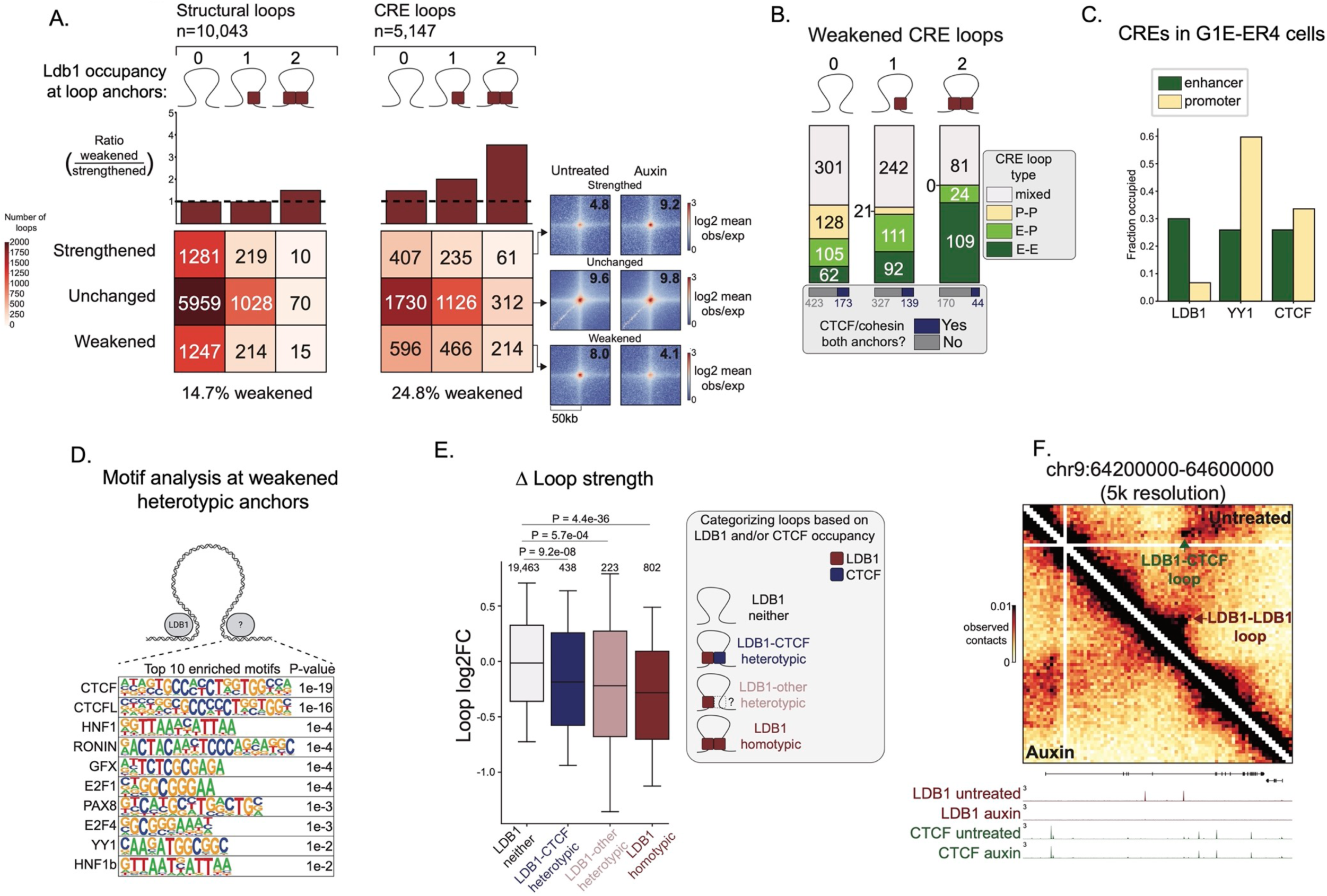
LDB1 mediates chromatin contacts between cis-regulatory elements. (A) Numbers of structural loops (left) and CRE loops (right) that are weakened (log2FC < −0.5), unchanged or strengthened (log2FC > 0.5) upon LDB1 depletion. Loops are stratified by LDB1 occupancy within anchors. (B) Distribution of CRE loop type for weakened CRE loops. Fraction of loops with RAD21/CTCF co-occupied peaks in both anchors (below). (C) Fraction of enhancers and promoters in G1E-ER4 cells occupied by LDB1 (left), YY1 (middle) and CTCF (right). (D) Schematic representing the motif analysis strategy for heterotypic loops and the top 10 most enriched motifs identified using HOMER known motif enrichment analysis. (E) Change in loop strength upon LDB1 depletion for loops categorized based on LDB1 and CTCF occupancy. Whiskers represent 10^th^ and 90^th^ percentiles; P-values calculated using a two-sided Mann-Whitney U test. (F) LDB1-dependent homotypic loop (red arrow) and LDB1-dependent heterotypic loop (green arrow).

We placed loops into 2 broad categories: 1) structural loops – CTCF/RAD21 at both anchors but CREs present only at one or no anchor, and 2) CRE loops-with CREs (enhancers or promoters) at both anchors. Upon LDB1 depletion 24.8% of CRE loops but only 14.7% of structural loops were weakened (Figure 1A). A similar number of structural loops were strengthened as were weakened. Additionally, only 16% of weakened structural loops contained LDB1 binding at one or both anchors, implying that rearrangements in structural loops may not be directly mediated by LDB1. In contrast, 53% of weakened CRE loops contained an LDB1 binding site in at least one anchor, suggesting that LDB1 preferentially maintains CRE loops.

To determine the type of CRE interactions dependent upon LDB1, we further stratified LDB1-dependent CRE loops into enhancer/enhancer (E-E), enhancer/promoter (E-P), promoter/promoter (P-P) or mixed (loops with both enhancer and promoter at a given anchor). Loops bound by LDB1 at both anchors were highly enriched for E-E interactions compared with loops that had LDB1 at only one anchor (Figure 1B), thus LDB1 is required for diverse CRE interactions. Importantly, most LDB1-dependent CRE loops lack CTCF/RAD21 co-bound sites at both anchors, suggesting that they formed independently of a CTCF-/cohesin mechanism.

Given that LDB1-dependent CRE loops with LDB1 present at both anchors are enriched for E-E interactions, we examined whether LDB1 preferentially binds to enhancers genome-wide. To do so, we intersected LDB1 ChIP-seq peaks with annotated CREs in G1E-ER4 cells and found that LDB1 occupancy favors enhancers over promoters (Figure 1C). To compare LDB1’s binding profile to other architectural factors, we integrated our CTCF ChIP-seq data. We also performed ChIP-seq for YY1 (a factor known to control subsets of enhancer/promoter loops). LDB1’s preference for enhancers is distinct from YY1 and CTCF that favor promoters (YY1) or have no preference (CTCF). These data suggest that LDB1 may have a unique role in enhancer connectivity and function through distinct mechanisms compared to other architectural proteins.

At the CAR2 locus, a heterotypic looping model has been proposed for LDB1 in which enhancer-bound LDB1 physically interacts with promoter-proximal CTCF^69^. To explore whether LDB1 engages broadly in heterotypic looping interactions with CTCF, we performed motif analysis at the LDB1-free loop anchors that are paired with an LDB1 occupied anchor of LDB1-dependent loops (using LDB1-dependent homotypic anchors as background regions to search for enriched motifs). CTCF was the most highly enriched motif, suggesting that LDB1 may broadly partner with CTCF to form loops (Figure 1D). To test the requirement of LDB1 for heterotypic loop configurations, we divided all loops into 4 categories based on LDB1 and CTCF occupancy: 1-“LDB1 neither loops” are not occupied by LDB1 at either anchor, 2-“LDB1-CTCF heterotypic loops” are occupied by LDB1 at one anchor and by CTCF at the opposite anchor (but do not have CTCF or LDB1 at both anchors), 3-“LDB1-other heterotypic loops are occupied by LDB1 at one anchor without CTCF at either anchor, 4-LDB1 homotypic loops” are occupied by LDB1 at both anchors. LDB1 homotypic loops were most sensitive to LDB1 depletion and LDB1 neither loops were least sensitive (Figure 1E). Moreover, both heterotypic categories (LDB1-CTCF and LDB1-other) were significantly more sensitive to LDB1 depletion compared to the LDB1 neither category. Examples of heterotypic and homotypic LDB1-dependent loops are shown in Figure 1F. These results are consistent with a broad role of LDB1 in connecting regulatory elements via homo-or heterotypic interactions.

Conventional loop calling may underestimate the number of CRE contacts if they are less frequent or if they encompass shorter genomic distances and are thus “overshadowed” by signal near the diagonal in the heat maps. To determine if there were additional LDB1-dependent loops that the Cooltools algorithm missed, we focused on LDB1 peaks that were not within identified loop anchors. We generated pairs of these LDB1 peaks (using a maximum distance between peaks of 500kb), quantified “loop” strengths for these paired sites using 2kb-binned Micro-C matrices, and filtered the list of paired sites to include those with a minimum observed/local expected value of 2 in the LDB1 replete Micro-C data set (a value representative of the weakest loops identified by Cooltools). We further filtered this list to include those with a CRE at both anchors and weakened upon LDB1 depletion (log2FC < −0.5). Using this strategy, we identified 660 additional putative LDB1-dependent CRE loops (Figure S1O). To test whether these putative LDB1-dependent CRE loops are missed by other loop calling algorithms, we identified loops using Mustache^81^ on the untreated Micro-C dataset. We used default parameters for 2kb, 5kb and 10kb resolution. Mustache was only able to identify 15% (99 of 660) of putative LDB1-dependent CRE loops. Together, these findings suggest that conventional loop calling using Micro-C data likely underestimates the total number of LDB1-dependent loops.

### LDB1 is acutely required for the nascent transcription of a subset of genes

To test the effects of acute LDB1 depletion on gene regulation, we performed TT-seq^82^ before/after 4 hours of LDB1 depletion to measure nascent transcription. The expression of 433 genes was reduced upon acute LDB1 depletion (log2FC <-1, padj <0.05) and of 480 genes was increased (log2FC >1, padj <0.05) (Figure 2A). Using a less stringent log2FC cutoff we identified an additional 1,064 genes that were downregulated (log2FC <-0.5, padj <0.05) and 818 genes that were upregulated (log2FC >0.5, padj <0.05) and characterized these as “weakly down/upregulated”. 7 genes with varying expression level changes were chosen and validated by primary transcript RT-qPCR (Figure S2B). These results demonstrate rapid LDB1-mediated changes in gene expression, suggesting its direct involvement in transcriptional regulation.

**Figure 2.**
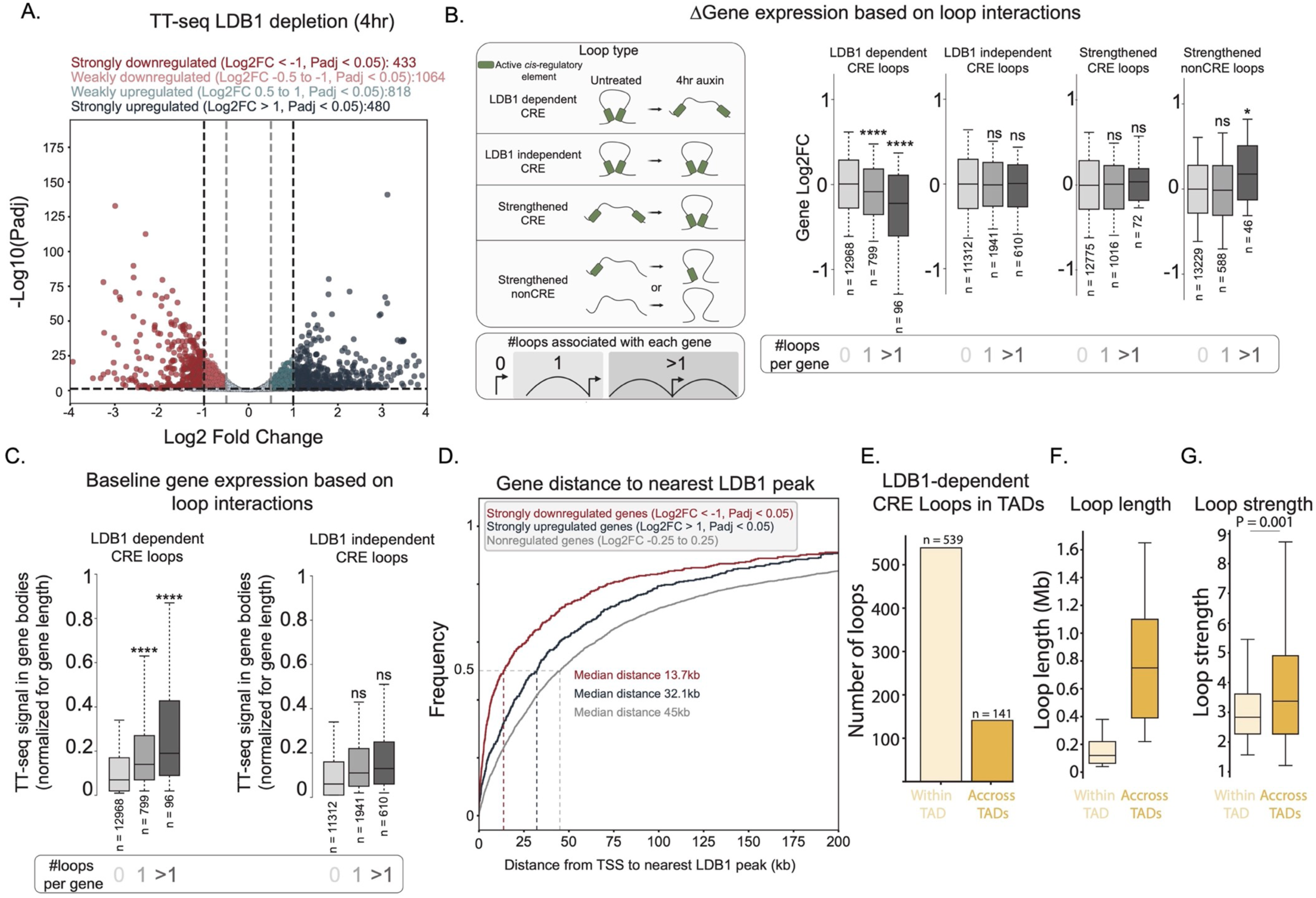
LDB1-dependent CRE loops are associated with transcription activation. (A) Gene expression changes measured by TT-seq upon LDB1 depletion (n=3). (B) Gene expression changes (TT-seq) for genes categorized by the number of loop anchors overlapping their TSS. Whiskers represent 10^th^ and 90^th^ percentiles; P-values calculated using a two-sided Mann-Whitney U test. *p<0.05, **p<0.01, ***p<0.001, ****p<0.0001. (C) Baseline gene expression measured by TT-seq. Genes categorized by the number of LDB1 dependent or independent CRE loops they interact with. (D) Cumulative frequency distributions for gene distance to nearest LDB1 ChIP-seq peak. (E) Numbers of inter-TAD vs intra-TAD LDB1-dependent CRE loops. (F) Loop lengths for LDB1-dependent inter-TAD and intra-TAD CRE loops. Whiskers represent 10^th^ and 90^th^ percentiles. (G) Loop strengths for LDB1-dependent inter-TAD and intra-TAD CRE loops. Loop strength calculated using 5k resolution.

LDB1-mediated enhancer interactions at the β-globin locus stimulate Pol2 recruitment to the β-globin promoter and subsequent early elongation^66^. To investigate whether LDB1 employs similar mechanisms to regulate transcription globally, we performed Pol2 ChIP-seq before/after 4 hours of LDB1 depletion. We measured Pol2 occupancy at transcription start site (TSS)-proximal regions (+/-750bp flanking the TSS) and transcription end sites (TES) (+1500bp downstream of TES). Additionally, we estimated the processivity of Pol2 by dividing the Pol2 ChIP-seq signal in TES regions by that in TSS regions for each gene (Pol2 TES/TSS). We focused our analysis on active genes by filtering for those enriched with the active H3K27ac mark at their TSS. Intriguingly, genes dependent upon LDB1 (downregulated upon LDB1 depletion) exhibited high Pol2 TES/TSS ratios at baseline compared to nonregulated or upregulated genes (Figure S2C) suggesting that LDB1 can drive high levels of transcription activation. Upon LDB1 depletion, downregulated genes showed a decrease in Pol2 occupancy at both their TSSs and TESs, and reduced Pol2 TES/TSS ratios. Conversely, upregulated genes showed an increase in Pol2 occupancy at both their TSSs and TESs and increased Pol2 TES/TSS ratios. Thus LDB1 likely modulates transcription by regulating Pol2 recruitment to promoters and may directly influence Pol2 elongation. However, the possibility remains that additional factors regulate Pol2 elongation after LDB1-mediated Pol2 recruitment.

### LDB1-dependent CRE loops are associated with transcription activation

To interrogate the relationship between LDB1’s role in looping and transcription regulation, we intersected the anchors of LDB1-dependent CRE loops with 1kb windows centered on TSSs. We focused on LDB1-dependent CRE loops at which LDB1 was detected at one or both anchors. Genes connected to LDB1-dependent CRE loops were more sensitive to LDB1 depletion than genes connected to LDB1 independent CRE loops (Figure 2B). Interestingly, genes overlapping multiple LDB1-dependent loop anchors were most sensitive to LDB1 depletion (Figure 2B). We performed the same analysis using our putative LDB1-dependent CRE loops from Figure S1O. Genes connected to LDB1-dependent putative CRE loops tended to be more sensitive to LDB1 depletion than genes that were not connected to LDB1-dependent putative CRE loops (Figure S2D). Hence, LDB1-mediated CRE connectivity is related to gene activation. We also measured the baseline gene expression (before auxin treatment) of genes that interacted with LDB1 dependent CRE loops. Genes connected to LDB1 dependent CRE loops tended to be expressed at higher levels compared to genes connected to LDB1 independent CRE loops (Figure 2C). This suggests that LDB1-mediated CRE interactions are associated with high levels of transcription activation.

To test whether loops strengthened in the absence of LDB1 were associated with transcription activation, we intersected the anchors of strengthened loops with 1kb windows centered on TSSs. We did so separately for strengthened loops with active CREs in both anchors and strengthened loops with active CREs in only one or no anchors (nonCRE loops) as upregulated genes may not have active H3K27ac prior to LDB1 depletion. Genes whose TSSs overlapped with multiple strengthened nonCRE loop anchors exhibited increased gene expression upon LDB1 depletion, however genes associated with strengthened CRE loops were not significantly changed (Figure 2B). Thus in some instances, upregulated genes can be explained by strengthened loops possibly resulting from aberrant interactions formed in the absence of LDB1.

Because of Micro-C resolution limits, short range LDB1 dependent loops are missing from our analyses. The shortest loops we could detect with Micro-C were 18kb long. To assess whether potential undetected short range LDB1 dependent loops may control gene expression, we measured the distance from LDB1 dependent genes (downregulated upon LDB1 depletion) to the nearest LDB1 binding site. As controls, we did the same for upregulated genes and genes that are not regulated by LDB1 (defined as those with log2FC values between −0.25 and 0.25). LDB1 was bound more proximally to downregulated genes than to upregulated or nonregulated genes. The median distance between LDB1 and downregulated genes was 13.7kb compared to 32.1kb for upregulated genes and 45kb for LDB1-insensitive genes (Figure 2D). We annotated the LDB1 binding sites relative to LDB1-dependent (downregulated) genes and found that LDB1 predominately occupies intronic (58%) and extragenic regions (30%) as opposed to promoter-proximal regions (8%) (defined as a 1kb window upstream of the TSS) (Figure S2E). Together, these findings suggest that LDB1 may mediate short range contacts to activate gene expression, many of which fall below our Micro-C loop detection limit. Additionally, LDB1 may often engage in heterotypic interactions to activate gene expression as many LDB1-dependent genes lack LDB1 occupancy at their promoters.

### LDB1 can regulate interactions across TAD boundaries

TADs are generally thought to constrain enhancer action; however, some enhancers can act across TAD boundaries^32,83–85^. To examine whether LDB1 regulatory influence can extend beyond TADs, we determined the number of LDB1-dependent CRE loops within TADs and those that crossed TAD boundaries. We identified TADs using the rGMAP^86^ algorithm and quantified loops with anchors within the same TAD or those with anchors in different TADs. While the majority of LDB1-dependent CRE loops reside within a given TAD, a considerable fraction crosses TAD boundaries (Figure 2E). Inter-TAD loops are substantially longer than intra-TAD loops (Figure 2F). Additionally, inter-TAD loops tended to be stronger (higher observed/locally-adjusted expected values) (Figure 2G) however they both exhibited the same fraction of homotypic/heterotypic LDB1 configurations and enhancer/enhancer vs enhancer/promoter interactions. These findings were corroborated using an independent TAD caller: HiTAD from TADLib^87,88^. Together, these data suggest that while LDB1 acts mostly within the confines of TADs it is also associated with inter-TAD interactions.

### LDB1 forms fine-scale looped networks at LDB1-dependent genes

LDB1 may mediate genomic contacts that escape detection by Micro-C (Figure 2D). Region-Capture Micro-C (RCMC) enhances the detection ability of Micro-C by capturing regions of interest prior to sequencing. We performed RCMC in LDB1-degron cells with/without 4 hours of auxin treatment. We used tiled capture probes to enrich for 5 distinct regions each ranging from 1-1.9 mb in length. Regions were chosen that harbored LDB1-dependent genes lacking associated LDB1-dependent Micro-C loops. We hypothesized that LDB1 may control small-scale looping interactions at these genes that were undetectable by genome-wide Micro-C.

RCMC uncovered a new layer of chromatin interactions that was undetectable by Micro-C (for a direct comparison between RCMC and Micro-C see Figure S3A). We used similar approaches to identify LDB1-dependent loops as we did for Micro-C with two adaptations specific for RCMC: 1-higher resolutions were applied to identify loops (500bp, 1kb, 2kb, and 5kb), 2-we relaxed loop calling parameters designed to merge nearby loops because RCMC can more reliably distinguish contacts in close proximity. Using this approach, we identified nearly seven times as many LDB1-dependent loops within captured regions (Figure 3A). RCMC highlights the connectivity of LDB1-driven contacts as many LDB1-dependent loops share anchors with each other. RCMC revealed that most LDB1 peaks within captured regions are affiliated with at least one weakened loop and over 40% of them are affiliated with more than 1 distinct weakened loop (Figure 3B). This contrasts with our Micro-C experiments which failed to detect most of these contacts. Similar to the Micro-C analysis, the presence of LDB1 at loop anchors is associated with the sensitivity of loops to LDB1 depletion (Figure S3B). In agreement with our Micro-C analysis, genes affiliated with multiple LDB1-dependent CRE loops (identified via RCMC) were most sensitive to LDB1 depletion (Figure S3C). Thus, the RCMC identified LDB1-dependent loops are linked to gene activation. Visually, LDB1-dependent loci seem to be part of LDB1-dependent multi looped networks. Examples of 5 LDB1-dependent genes (Zfpm1, Uba7, Myc, Cbfa2t3, and Bcl2l1) are shown in Figures 3C and S3D-E. Intriguingly, at the Zfpm1 and Uba7 loci, LDB1 forms looped networks that are flanked by invariant CTCF/cohesin bound loops, whereas at the Cbfa2t3 locus and Myc proximal region, LDB1-dependent contacts share an anchor with an encompassing CTCF/cohesin-occupied loop that is also sensitive to LDB1 depletion. At some sites (such as the Myc proximal region) LDB1 degradation reduces cohesin occupancy, yet at others (such as the Zfpm1 locus) LDB1 is dispensable for cohesin binding. We explore the requirement of LDB1 for cohesin occupancy genome-wide below in Figure 4. Together, these results hint at potentially cohesin-independent but also partially cohesin-dependent roles for LDB1 loop formation that may be locus specific.

**Figure 3.**
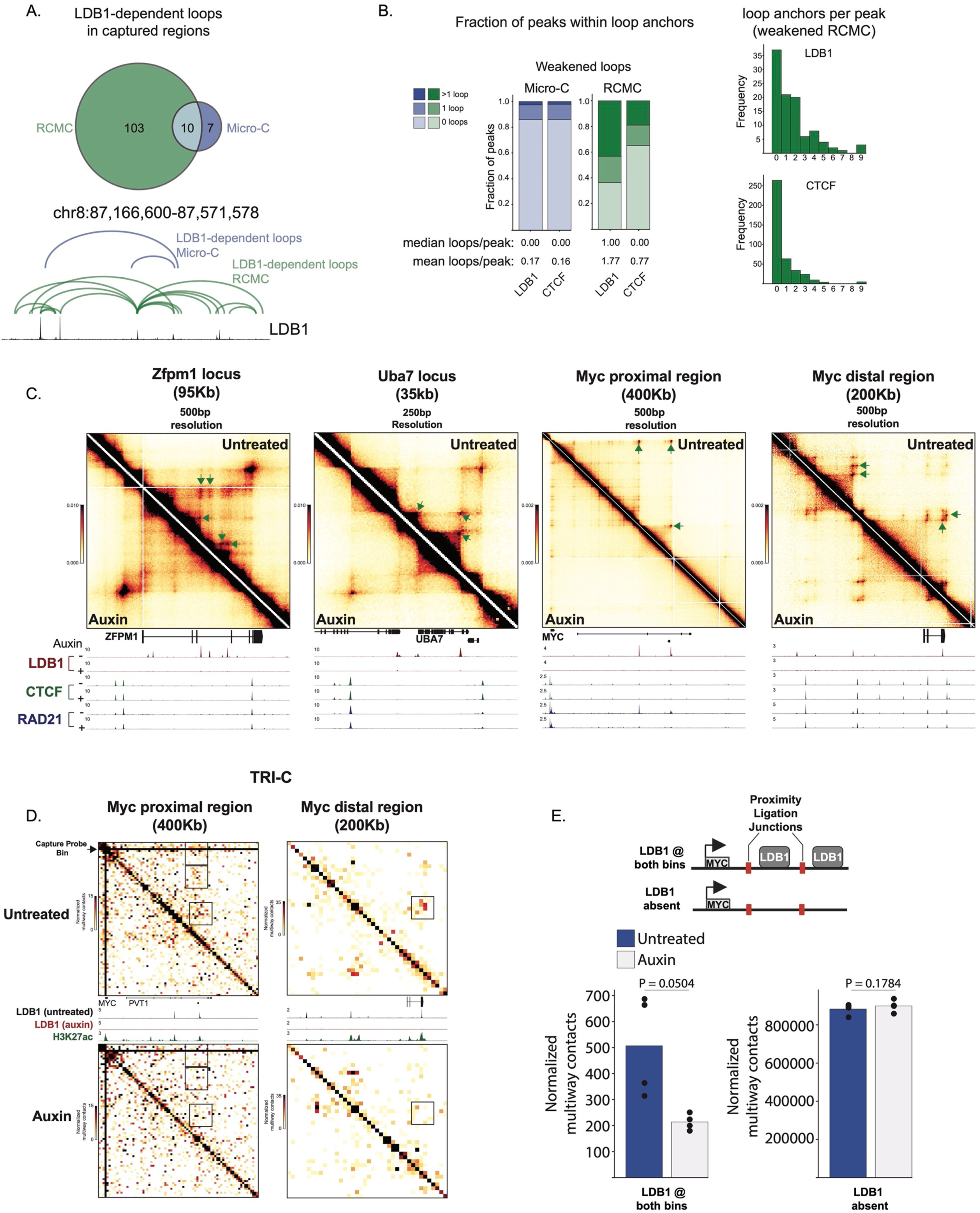
LDB1 forms fine-scale looped networks at LDB1-dependent genes. (A) Numbers of LDB1-dependent loops detected by Micro-C or RCMC. ChIP-seq tracks for LDB1 are shown in black. (B) Proportions of LDB1 or CTCF ChIP-seq peaks overlapping weakened loop anchors identified by Micro-C (blue) or RCMC (green). For overlaps with RCMC, only peaks within captured regions are considered. Histograms (right) showing the number of LDB1 or CTCF peaks that overlap with increasing numbers of weakened loop anchors identified by RCMC. (C) Examples of LDB1-dependent looped networks. Green arrows indicate LDB1-dependent loops. (D) 5k resolution TRI-C contact maps for MYC proximal and distal regions. Contacts represent multi-way interactions involving the MYC promoter. Capture probe bin indicated by black arrow. (E) Multiway contacts with the MYC promoter and bins occupied by LDB1 or unoccupied by LDB1. Dots represent normalized multiway contacts for each biological replicate. P values calculated using paired t-test.

**Figure 4.**
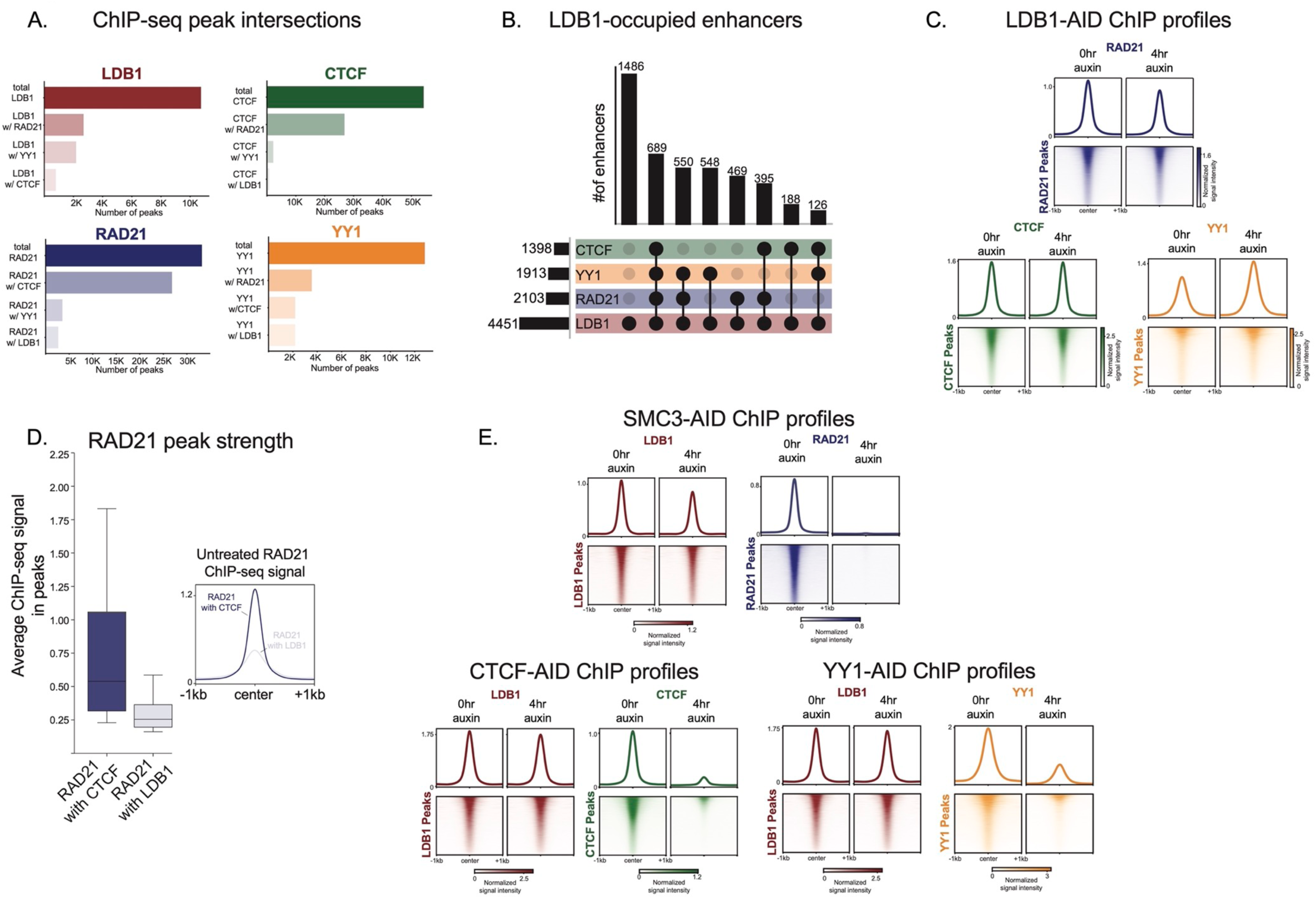
LDB1 occupancy is mutually independent of YY1, CTCF and cohesin at most locations. (A) ChIP-seq peak intersections between LDB1, CTCF, RAD21 and YY1. (B) LDB1-occupied enhancer elements that are occupied by cohesin (RAD21), YY1, or CTCF. (C) ChIP-seq profiles in LDB1-AID cells for RAD21, CTCF and YY1 before/after LDB1 depletion. Heatmaps and profiles are shown for peaks identified for each factor in the LDB1 replete condition. (D) RAD21 ChIP-seq signal at RAD21 ChIP-seq peaks overlapping CTCF peaks or LDB1 peaks. Whiskers are 10^th^ and 90^th^ percentile. (E) ChIP-seq profiles in SMC3-AID, CTCF-AID, and YY1-AID cells before/after 4hr auxin treatment.

A substantial fraction of LDB1 loop anchors engage in multiple contacts, raising the question whether they occur in a mutually exclusive manner or whether some are capable of forming simultaneous multi-way intra-allelic contacts to form enhancer ehubs^89^. To this end, we performed Tri-C^78^ which enables detection of multiway contacts between loci of interest. We focused on the Myc locus because LDB1-occupied enhancers are relatively widely spaced, enabling detection of simultaneous contacts with moderate resolution. We used a capture probe proximal to the Myc TSS to enrich for contacts with the Myc promoter region. Using the Capcruncher^90^ analysis pipeline, we filtered for read fragments that contain the capture site and at least two additional fragments separated by a restriction enzyme recognition site (NLAIII) and plotted the contact frequencies of only these filtered fragments as a heatmap. Thus, reads on the heatmap represent multiway interactions between the Myc promoter and at least two additional sites. We binned our contact matrix at 5kb resolution and found that simultaneous, multiway contacts were enriched at LDB1-binding sites (both in the proximal and distal clusters; indicated by black squares), and that these contacts were diminished upon auxin treatment (Figure 3D). We quantified contacts involving the Myc promoter and found that LDB1 depletion resulted in diminished simultaneous contacts between the Myc promoter and distinct regions bound by LDB1 (Figure 3E). While comparing absolute frequencies of multi-way vs two-way interactions is challenging, our results support the idea that in principle simultaneous LDB1-dependent multi-way contacts can form among LDB1-occupied sites.

### LDB1 occupies distinct genomic loci relative to YY1, CTCF and cohesin

Structural loops formed by the CTCF/cohesin machinery can support or interfere with E-P loop formation^91,92^. Moreover, the cohesin-mediated chromatid extrusion process may increase the likelihood of an E-P encounter. Separately, YY1 has been proposed as a general E-P looping factor genome-wide^39^ although the extent to which YY1 regulates E-P interactions globally is debated^32^. To explore the mechanisms through which LDB1 forges CRE loops, we began by testing any functional relationship between LDB1 and other well-studied architectural factors (CTCF, cohesin and YY1).

We began to examine relationships among LDB1, CTCF, cohesin, and YY1 by comparing the ChIP-seq profiles in cells carrying the LDB1-degron fusion protein. LDB1 predominantly binds in a manner mutually exclusive to that of the other factors (Figure 4A). 75% of LDB1 peaks did not intersect with RAD21 peaks, 93% of LDB1 peaks did not intersect with CTCF peaks, and 80% of LDB1 peaks did not intersect with YY1 peaks. To explore whether CTCF, cohesin or YY1 influence LDB1’s effect on enhancers, we assessed their presence across LDB1-bound enhancer elements. We found that LDB1 often binds to enhancers in the absence of the other architectural factors, suggesting that LDB1 may not rely on CTCF, cohesin, or YY1 for its function (Figure 4B).

### YY1, CTCF and cohesin occupancy is not regulated by LDB1 at most locations

Any interpretation of LDB1 loss-of-function experiments must consider that LDB1 may affect the chromatin occupancy of other factors, for example via protein-protein interactions, via chromatin binding cooperativity, or in the case of cohesin, via stalling loop extrusion. To test the influence of LDB1 on the binding of other architectural factors, we measured their genomic occupancy profiles following LDB1 depletion. Globally, YY1, CTCF, and RAD21 occupancy was largely unaffected by LDB1 depletion (Figure 4C). However, we observed a modest reduction in RAD21 occupancy specifically at LDB1 co-occupied sites (Figure S4A). However, at these sites, RAD21 occupancy was much lower than at CTCF/RAD21 co-bound sites (Figure 4D), suggesting that LDB1 is, if at all, an ineffective cohesin extrusion blocker. More likely, loss of LDB1 may directly or indirectly affect cohesin loading at a subset of sites.

Since LDB1 chromatin occupancy occurs predominantly at enhancers (Figure 1C), we explored whether LDB1 dependent cohesin enrichment also occurs at enhancers. First we quantified changes in ChIP-seq signal at each RAD21 peak and identified only 2,284 out of 33,204 (7%) peaks to be weakened upon LDB1 depletion (by at least 50%). Of these, 906 (40%) were located at LDB1-occupied enhancers. Conversely, only 805/4,451 (18%) of LDB1-occupied enhancers were associated with RAD21 peaks that were modulated by LDB1 (Figure S4B-C). Hence, LDB1 predominantly influences enhancer connectivity independently of cohesin levels.

### LDB1-dependent looping is uncoupled from YY1, CTCF and cohesin occupancy

While LDB1 did not substantially influence YY1, CTCF or cohesin occupancy globally, the possibility remained that these factors may be diminished specifically at LDB1-dependent loop anchors. To assess the influence of YY1, CTCF and cohesin reduction upon LDB1 depletion on chromatin looping, we determined the number of weakened CRE loops with diminished (by at last 50%) YY1, CTCF or RAD21 peaks at their anchors using diminished LDB1 peaks as a control. Most weakened CRE loops did not harbor reduced YY1, CTCF or cohesin sites. Importantly, weakened LDB1 peaks were enriched at weakened CRE loop anchors relative to the other factors (Figure S4D). Thus, LDB1 dependent loops are unlikely to be significantly influenced by changes in YY1, CTCF or cohesin occupancy.

We next investigated whether strengthened loops upon LDB1 depletion were influenced by positive changes in YY1, CTCF or cohesin occupancy. To do so, we measured the number of strengthened loops occupied by strengthened (by at least 50%) YY1, CTCF, and RAD21 peaks. We included strengthened peaks exclusively identified in the 4hr auxin condition to determine if any de-novo peaks contributed to strengthened loops in the absence of LDB1. Very few strengthened CRE loops harbored strengthened RAD21 (10) or CTCF (4) sites in either anchor (Figure S4E). Conversely, 275/703 strengthened CRE loops were occupied by strengthened YY1 sites in one anchor and 39/703 were occupied by strengthened YY1 peaks in both anchors. YY1 is present at many (∼60%, Figure 1C) active promoters in G1E-ER4 cells, thus to determine if strengthened YY1 peaks were specifically enriched at strengthened loops we also determined their presence at weakened CRE loop anchors. We found that similar fractions of weakened loops were occupied by strengthened YY1 peaks suggesting that YY1 may simply be present at many active CREs and does not necessarily influence changes in chromatin looping upon LDB1 depletion (Figure S4F).

In sum, a substantial fraction of LDB1’s architectural functions may be uncoupled from those involving CTCF, cohesin, and YY1. However, the possibilities remain that a subset of LDB1-dependent contacts may be mediated by heterotypic protein complexes such as LDB1-CTCF^69^, and that the process of loop extrusion aids in the formation of LDB1 anchored loops.

### YY1, CTCF, and Cohesin do not influence LDB1 occupancy

Cohesin can influence transcription factor binding^32,93–95^. Hence, looped contacts lost upon cohesin depletion may be caused by reduced occupancy of architectural transcription factors, independently of the cohesin loop extrusion process. We therefore examined whether the cohesin dependency of a subset of loops may be explained by loss of LDB1 binding. We carried out LDB1 ChIP-seq in a G1E-ER4 line in which the SMC3 subunit of cohesin was tagged with an AID domain (Zhao et. al., in press) before and after exposure to auxin for 4 hours. Only 374 LDB1 peaks (3.5%) exhibited a ≥ 50% reduction in LDB1 ChIP-seq signal strength, indicating that SMC3 loss had little effect on LDB1 chromatin occupancy (Figure 4E, Figure S4C). As a control, 95% of RAD21 peaks were diminished by ≥50% (Figure 4E, Figure S4C). These results support the idea that LDB1 genomic occupancy is not substantially influenced by cohesin within the measured time frame, and that the majority of cohesin dependent loops cannot be explained by changes in LDB1 occupancy.

We next tested whether LDB1 occupancy was influenced by CTCF or YY1 by performing LDB1 ChIP-seq in G1E-ER4 cells in which CTCF^26^ or YY1 (Lam et. al., under review) was tagged with an AID moiety. LDB1 occupancy was not affected by loss of either factor supporting the idea that LDB1 occupancy is independent of CTCF and YY1 (Figure 4E, Figure S4C).

### LDB1 dependent loops can form in the absence of cohesin

LDB1 occupancy is uncoupled from that of CTCF/cohesin at most locations. Yet, structural loops mediated by CTCF/cohesin or the loop extrusion process itself may influence LDB1 dependent CRE loops. To this end, we analyzed Hi-C data generated in SMC3-AID G1E-ER4 cells treated with auxin for 4 hours (Zhao et. al., in press). Perturbing cohesin via SMC3 degradation allowed us to simultaneously test the influence that structural loops and the process of loop extrusion have on LDB1 dependent CRE loops. Focusing on LDB1-dependent CRE loops at which LDB1 was detected at one or both anchors we calculated the change in loop strength and assigned loops to one of two categories: LDB1-dependent and SMC3-independent (LDB1-AID log2FC < −0.5, SMC3-AID log2FC > −0.5), and loops dependent on both (LDB1-AID log2FC < −0.5, SMC3-AID log2FC < −0.5). Using this binary categorization, 70% of LDB1-dependent loops were unaffected by SMC3 depletion (Figure 5A). Cohesin-independent loops tended to involve stronger enhancers (as measured by the active mark H3K27ac^96^) (Figure 5B). These results suggest that the majority of LDB1-dependent loops form independently of cohesin.

**Figure 5.**
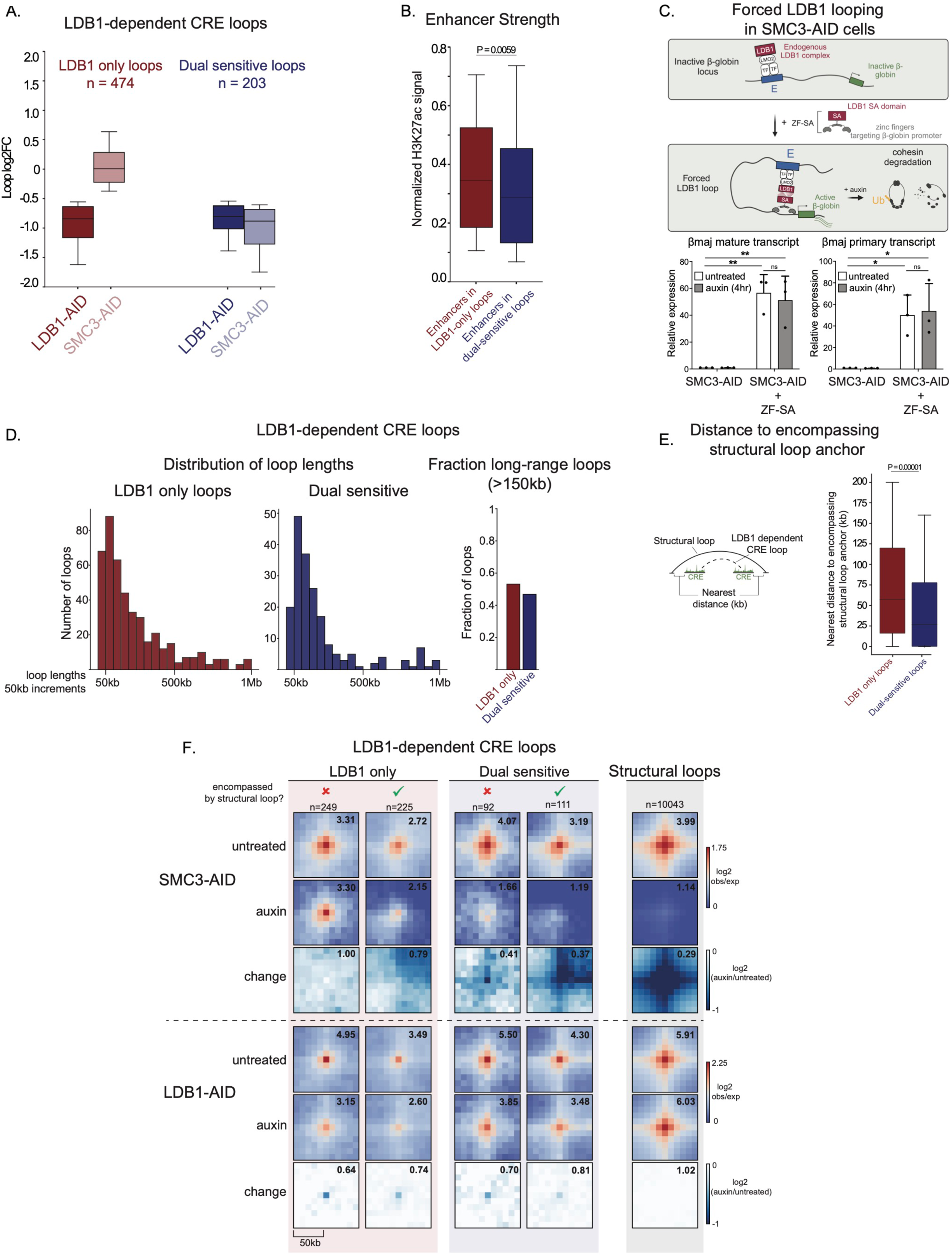
LDB1 Can Function in the absence of cohesin. (A) Change in loop strength for LDB1-dependent CRE loops in response to LDB1 depletion (darker colors) or SMC3 depletion (lighter colors). Loops are categorized as LDB1 only loops (red) or dual sensitive loops (blue). (B) H3K27ac ChIP-seq signal at enhancers within LDB1-only loop anchors or dually-sensitive loop anchors. Only mutually exclusive enhancer elements between the two sets are considered. (C) Relative RNA levels for β-globin measured by RT-qPCR in SMC3-AID cells −/+ ZF-SA and −/+ auxin (4hr). P-values calculated using One-way ANOVA. *p < 0.05, **p < 0.01. (D) Lengths of LDB1 only and LDB1/cohesin dual sensitive loops. (E) Distance to encompassing structural loop anchors for LDB1-only loops and dually sensitive loops. Only loops with an encompassing structural loop are shown. (F) APA plots for LDB1-dependent CRE loops stratified by their response to SMC3 depletion and whether they are encompassed by a structural loop. Numbers represent raw center pixel values.

While globally, cohesin depletion had little effect on LDB1 chromatin occupancy (see above) it remained possible that cohesin loss diminished LDB1 chromatin occupancy specifically at anchors of LDB1/cohesin-dependent CRE loops. To test this possibility, we measured LDB1 ChIP-seq peak signals at the anchors of loops dependent on both LDB1 and cohesin. LDB1 ChIP-seq signal was not substantially altered at LDB1/cohesin-dependent loop anchors (Figure S5A). Thus, loss of LDB1 occupancy does not explain the cohesin requirement for a subset of LDB1 dependent loops. Conversely, LDB1 does not influence cohesin occupancy at most locations, but it remained possible that LDB1 loss diminished cohesin occupancy specifically at the anchors of LDB1/cohesin-dependent CRE loops. To test this possibility, we quantified the number of weakened RAD21 ChIP-seq peaks (upon LDB1 depletion) present in LDB1/cohesin dually dependent CRE loop anchors. Only 5 such loops (2.5%) had weakened RAD21 peaks in both anchors and 52 (25.6%) had weakened RAD21 peaks in one anchor (Figure S5A). Thus, loss of cohesin occupancy does not explain the cohesin requirement for a subset of LDB1 dependent loops.

### Cohesin is dispensable for an engineered LDB1-dependent chromatin loop

Studies that assess endogenous CRE loops for their dependence on LDB1 and cohesin may be confounded by the general complexities of CREs. For example, changes in LDB1 or cohesin levels may impact other enhancer or promoter-bound factors that contribute to long range chromatin contacts. We therefore employed a defined system, in which a chromatin loop can be engineered at the murine β-globin locus via targeted LDB1 recruitment^66,67^. In this system, an artificial zinc finger (ZF) protein that binds to the β-globin promoter is fused to LDB1 or its self-association (SA) domain and introduced into G1E erythroid cells lacking transcription factor GATA1. In the absence of GATA1, β-globin promoter-enhancer contacts are rare. However, expression of ZF-LDB1 or ZF-SA establishes strong E-P contacts^66,67^ and activates β-globin transcription in a manner dependent on the enhancer. To examine whether cohesin is required for LDB1 function during this process, we introduced ZF-SA into undifferentiated SMC3-AID G1E-ER4 cells, treated cells with auxin, and measured β-globin expression via RT-qPCR. As expected, ZF-SA strongly induced β-globin transcription (∼50 fold). Importantly, depletion of cohesin for 4 hours had no effect on β-globin transcription activation, consistent with the dispensability of cohesin for LDB1 looping function in this system (Figure 5C).

### LDB1 can mediate long-range CRE interactions independent of cohesin

Previous reports suggest that cohesin may be required for long-range CRE interactions^28,29^. To test the requirement of cohesin for long-range LDB1 interactions, we measured the length of loops exclusively dependent on LDB1 and those dependent on both LDB1 and cohesin by measuring the distances between each of their respective anchors. Both LDB1 only and LDB1/cohesin dually dependent interactions spanned a wide range of distances, with many extending beyond 150kb (Figure 5D). Thus LDB1 can forge long-range contacts independent of cohesin.

### LDB1 dependent loops can be supported by structural loops or the process of loop extrusion itself

The effect of cohesin on 30% of LDB1 dependent loops may be due to the extrusion process itself or due to encompassing supportive structural CTCF/cohesin loops^26,54,97^. To distinguish between these possibilities, we categorized LDB1-dependent loops into two groups: those encompassed by a structural loop and those that are not. We then measured the distance of each loop to its encompassing structural loop. Similar fractions of LDB1 only dependent loops and dually dependent loops were encompassed by a structural loop (47.5% and 54.7% respectively). However, for those encompassed by a structural loop, dually dependent loops were located significantly closer to an encompassing structural loop anchor than were loops exclusively dependent on LDB1 (Figure 5E). Hence, LDB1 dependent loops benefited from structural loops when in close juxtaposition. Aggregate Peak Analysis (APA) plots showing the average contact frequencies for all LDB1 dependent CRE loop subtypes before/after either LDB1 or SMC3 depletion are shown in Figure 5F. These data suggest that nearby CTCF/cohesin-bound structural loops may facilitate a subset of LDB1 dependent loops. However, since roughly half of dually dependent loops are not encompassed by a structural loop, the influence of structural loops does not completely account for cohesin’s impact on LDB1 dependent loop formation. Thus, the extrusion process itself may separately facilitate the formation of a subset of LDB1 dependent loops.

To independently assess the role of structural loops on LDB1-dependent contacts we analyzed published Hi-C data from CTCF-AID G1E-ER4 cells^26^. These data sets were generated from CTCF-depleted cells transitioning from mitosis to G1-phase, providing the added advantage of testing structural loop requirements for the establishment (as opposed to maintenance) of LDB1-dependent loops during G1-phase entry. The majority of LDB1-dependent loops formed normally in the absence of CTCF depletion (Figure S5B). LDB1 loops that were not influenced by CTCF tended to be more distal to encompassing structural loops and included stronger enhancer elements than did dually dependent loops (Figure S5C-D). CTCF depletion did not affect LDB1 occupancy at CTCF/LDB1 dually-dependent loop anchors (Figure S5A). Leveraging the CTCF-AID and SMC3-AID degron systems, we were able to distinguish between the impacts of structural loops as opposed to the loop extrusion process itself on LDB1 dependent loops. By comparing the LDB1/SMC3 dually dependent loops to the LDB1/CTCF dually dependent loops, we found that 37% of LDB1/SMC3 dually dependent loops were also sensitive to CTCF depletion (Figure S5A). Thus, in cases where cohesin facilitates LDB1 dependent loops, it predominately does so through active extrusion, and in a minority of cases can do so through the formation of structural loops where cohesin is stalled by CTCF.

### LDB1 regulates distinct CRE loops compared to YY1

YY1 has been proposed to function as a global connector of CRE loops^39^, yet many CRE loops remain intact following acute YY1 depletion^32^. Therefore, alternative factors may control CRE loops in a manner distinct to YY1. Because LDB1 has minimal genomic overlap with YY1 and preferentially binds enhancers (as opposed to YY1 which preferentially binds promoters), we suspected that LDB1 may forge regulatory loops through distinct mechanisms and may even control different subsets of CRE loops. To this end, we utilized Micro-C data from YY1-AID G1E-ER4 cells (Lam et. al., under review) and found that 90% of LDB1-dependent CRE loops persisted during the acute absence of YY1 (Figure S5E). As opposed to our findings using the CTCF-AID and SMC3-AID systems, neither enhancer strength nor distance to structural loop anchors were predictive of whether a loop was exclusively dependent upon LDB1 or dependent on both LDB1 and YY1 (Figure SF-G). These findings demonstrate that LDB1 regulates distinct CRE loops compared to YY1 providing an explanation for why acute YY1 depletion does not result in global loss of E-P loops.

### LDB1 chromatin occupancy is associated with loop establishment during G1-phase entry

Mitosis is an interval during which gene expression, transcription factor occupancy and loops are temporarily disrupted^98–101^. The study of cells transitioning into G1-phase presents a powerful opportunity to test the correlation between LDB1 chromatin occupancy, loop formation and gene expression. We performed ChIP-seq for LDB1 in highly purified cell populations at closely spaced timepoints including prometaphase, ana/telophase, early G1, mid G1 and late G1 and compared signals at LDB1 peaks identified in asynchronous cells. LDB1 is essentially undetectable in prometaphase and gradually strengthens through mid G1 (Figure 6A). We integrated our ChIP-seq data with published Hi-C data collected from G1E-ER4 cells at the same cell cycle stages^36^. Measuring the average Hi-C signal of LDB1-dependent CRE loops (as defined using asynchronous cells) and the average ChIP-seq signal for LDB1 at loop anchors of LDB1-dependent loops, revealed that LDB1 occupancy at loop anchors is associated with the re-formation of loops during mitotic exit (Figure 6B).

**Figure 6.**
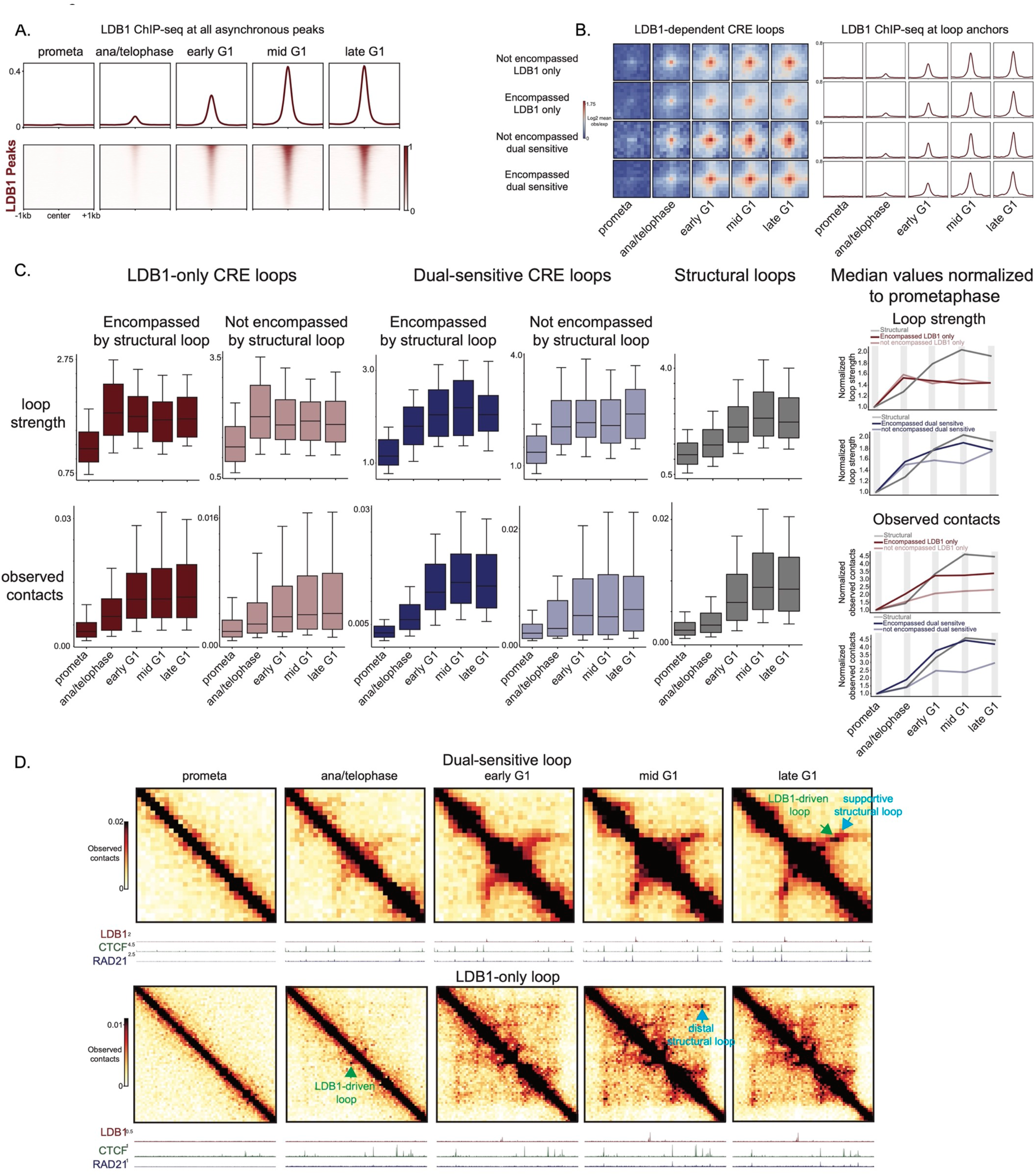
LDB1 chromatin occupancy correlates with loop establishment during G1-phase entry. (A) ChIP-seq profiles for LDB1 at each cell cycle stage at all LDB1 peaks identified in asynchronous cells. (B) APA plots from 10k resolution Hi-C data at each cell cycle stage for each category of LDB1-dependent CRE loops. Average ChIP-seq profiles are shown for each loop type for LDB1 peaks within loop anchors. (C) Loop strength (top) and observed contacts between loop anchors (bottom) for each category of LDB1-dependent CRE loops and for structural loops at each cell cycle stage. Median loop strength and observed contacts normalized to prometaphase are shown for each loop category (right). (D) Examples of an LDB1/cohesin dually sensitive loop (top) and LDB1-only loop (bottom). Green arrow indicates the LDB1-dependent loop, blue arrow indicates an encompassing structural loop.

Previous studies found that during G1-phase entry, CRE loops can form before cohesin-driven structural loops, and uncoupled from TAD formation, implying that CRE loops may not require support from structural loops^26,36^. To investigate the dynamics of LDB1-dependent loop formation in the context of structural loops, we again stratified LDB1-dependent CRE loops into LDB1-only and LDB1/cohesin dually dependent loops. We subdivided each group into those encompassed by a structural loop and those that are not. For each loop type, we measured loop strength as we did previously by quantifying the observed contacts/locally-adjusted expected value between loop anchors for each loop at each cell cycle stage. Additionally, we simply measured the observed contacts between loop anchors at each stage. We found that LDB1-only loops were established more rapidly during mitotic exit relative to their local background compared to dually-dependent loops (Figure 6C). LDB1-only loops reached maximum loop strength values in ana/telophase while dual-sensitive loop dynamics more closely mimicked those of structural loops with a gradual increase in loop strength through mid/late G1. Both LDB1-only and dually-dependent loops exhibited a gradual increase in the observed contacts between anchors, however for LDB1 only loops the increase in contact frequency was modest after early G1. Thus, the absolute contact frequency for all loops is gradually increased during mitotic exit; however, LDB1-only loops are established more rapidly relative to their local background than dually-dependent loops. Examples showing the formation of a dually-dependent and LDB1-only loop are shown in Figure 6D. These results support the idea that LDB1-only loops are not only maintained in the absence of cohesin, but may be established independently of cohesin; whereas, dual-sensitive loops rely on cohesin-mediated loop extrusion for establishment and maintenance.

## DISCUCSSION

Only a select few nuclear factors have been studied for a direct/proximal role in CRE connectivity. Using a 4 hr LDB1 depletion scheme, we identified a widespread role for LDB1 in organizing CRE interactions and maintaining transcription regulation. We further discovered that LDB1 can support complex E-P networks (also termed hubs^43,89,102–104^). 1-LDB1 mediates E-E as well as E-P loops, 2-LDB1-dependent loops display a high level of connectivity and often share anchors with each other, 3-Shared contacts can occur simultaneously based on Tri-C experiments. Such hubs may convey high level transcriptional output. Although we lack estimates of the number of LDB1 controlled hubs genome wide, we speculate that they are quite common as suggested by high RNA levels of genes connected to multiple LDB1 dependent loops.

Mechanistically, we found no evidence that LDB1 functions as a loop extrusion blocker analogous to CTCF. 1-LDB1 lacks co-occupancy with cohesin, 2-in the minority of cases where cohesin does occupy an LDB1 site, cohesin binding is much weaker compared to CTCF sites, 3-most LDB1 dependent CRE loops lack CTCF/cohesin occupancy at both anchors, 4-the majority of LDB1 dependent loops are maintained upon acute cohesin depletion, including a specifically engineered loop formed by targeted LDB1 tethering. These findings suggest that these loops require neither the support of structural loops/TADs nor the process of cohesin extrusion per se in order to be formed. Moreover, while previous reports suggest that some long-range E-P contacts require cohesin^28,29^, we find numerous LDB1-dependent contacts (>150kb) that are cohesin independent. Lastly, using a degron approach, we also ruled out YY1, another factor with presumed wide-spread roles in E-P connectivity as a major force in LDB1 dependent looping.

However, a subset of LDB1 dependent loops is supported by CTCF/cohesin-anchored structural loops if their respective anchors are in close proximity. To uncouple structural loop support from a potential role of the loop extrusion process per se, we took advantage of our ability to selectively perturb CTCF and cohesin independently of each other, which revealed LDB1 loops that are dependent on cohesin but independent of CTCF. This supports cohesin extrusion as an additional mechanism to promote LDB1-anchored contacts. These results are buttressed by the dynamics of LDB1 loop formation in cells exiting mitosis. It is possible that the positive, negative or neutral influence of structural loops and cohesin-driven loop extrusion on LDB1 loops is a general reflection of the various ways by which CTCF and cohesin modulate CRE contacts.

We uncovered LDB1 dependent loops with LDB1 on one (heterotypic) or both (homotypic) anchors, suggesting that LDB1 can partner with non-self proteins. Homotypic LDB1 loops tended to involve E-E interactions whereas heterotypic LDB1 loops tended to involve E-P interactions. While CTCF is present at the opposite anchors of some heterotypic LDB1 loops and may function as a direct partner^69^, a substantial number lack CTCF binding. Hence, LDB1 may engage with other yet to be characterized partners. While demonstrating a direct role for protein multimerization or heterodimerization in loop formation in vivo is challenging, the most parsimonious working model is that LDB1 forms oligomers likely involving additional partners to form multimolecular assemblies that connect CREs.

LDB1 dependent loops were generally associated with transcription activation, yet a number of genes reliant upon LDB1 lacked Micro-C-detectable loops involving their promoters. Using RCMC, we uncovered new LDB1-dependent short-range contacts that escaped detection by Micro-C, suggesting that the number of functionally important LDB1-anchored loops is likely much higher than what is observed with genome-wide Hi-C/Micro-C.

A subset of genes also exhibited increased expression upon LDB1 depletion as measured by TT-seq. The mechanism may reflect direct repression of these genes by LDB1 as suggested in prior studies^105–107^. However, our data additionally suggest that in the absence of LDB1, new loops are formed to increase gene transcription. We propose that by its ability to forge connectivity networks, LDB1 also prevents illegitimate regulatory contacts.

To distinguish the role of LDB1 during establishment vs maintenance of CRE loops, we measured the formation kinetics of LDB1 dependent loops during the mitosis-G1-phase transition. LDB1 is evicted from mitotic chromatin, and its rapid re-binding was associated with loop re-formation. However, maximal loop intensities preceded peak LDB1 binding intensities. Possible explanations are 1. A non-linear relationship between LDB1 occupancy and loop formation, such as threshold effects. Also in asynchronously growing cells, LDB1 chip seq peak size and loop strength were not correlated (not shown). 2. The strong early appearance of LDB1 anchored loops is apparent on a background of few chromatin contacts and a virtual absence of domains and TADs, in other words low background signal against which focal CRE loops are quantified (observed/expected). Further gains in CRE loop strength will appear blunted upon gains in surrounding local interactions.

An additional informative observation derived from the cell cycle studies is that LDB1 dependent, cohesin independent loops can be established quickly and prior to structural loops. This lends further support to the idea that LDB1-dependent loops can not only persist, but also be established independent of cohesin/CTCF. These results are also consistent with our previous findings that general CRE connectivity can be established prior to and/or independently of cohesin and CTCF^26,36^.

In sum, by leveraging multiple degron systems (LDB1, cohesin, CTCF, YY1), cell cycle dynamics, and an engineered loop, our findings establish LDB1 as a major genome wide driver of CRE connectivity. This includes its ability to organize CRE hubs that are associated with high levels of transcription. CTCF, cohesin and YY1 may influence LDB1 connectivity in select circumstances but LDB1 can function in their absence, likely via homotypic and heterotypic protein complexes.

## SUPPLEMENTAL FIGURE LEGENDS

**Figure S1. Auxin-inducible degron system for LDB1, related to Figure 1**

(A) Schematic of LDB1-AID degron system.

(B) Western blot in whole cell lysates for parental G1E-ER4 cells and two LDB1-AID subclones in untreated and 4 hour auxin-treated conditions. GAPDH is shown as a loading control. Asterisks indicate nonspecific bands.

(C) Flow cytometry histograms for mCherry signal in asynchronous LDB1-AID cells upon auxin treatment. Flow cytometry histograms are representative of two independent experiments.

(D) Heat maps showing LDB1 ChIP-seq signal at all LDB1 peaks identified in the untreated condition.

(E) Bar plots showing relative nascent RNA levels for β-globin and Gypa measured by RT-qPCR normalized to Actin. RNA was extracted from parental G1E-ER4 cells and two LDB1-AID subclones. RNA extractions were performed under the following treatment conditions: without induction, after 24 hours of induction solely with estradiol, and after 24 hours of induction with estradiol in combination with a simultaneous auxin treatment. Bar graphs are representative of two independent experiments; dots represent technical replicates.

(F) Pearson correlation between parental G1E-ER4 cells and LDB1-AID clonal lines based on TPM values for all genes with TPM >1 (RNA-seq).

(G) Gene expression in G1E-ER4 parental cells and two LDB1-AID clones for genes categorized in parental G1E-ER4 cells (RNA-seq).

(H) Gene expression for genes located near LDB1 ChIP-seq peaks (within 50kb) (RNA-seq).

(I) Gene expression for LDB1 erythroid targets in parental G1E-ER4 cells and two LDB1-AID clones.

(J) APA plots for all Micro-C samples (1k resolution) performed in LDB1-AID cells. Plots shown for all weakened CRE loops, unchanged CRE loops, and strengthened CRE loops upon LDB1 depletion. Heatmap showing Pearson correlation among all Micro-C samples, based on eigenvector 1 of 100kb bins.

(K) Saddle plots showing compartment strength in LDB1-AID cells in untreated and 4 hour auxin-treated conditions.

(L) Insulation scores at TAD boundaries in LDB1-AID cells in untreated and 4 hour auxin-treated conditions. TADs identified using rGMAP on 10k resolution Micro-C matrices. Insulation scores calculated using a 120kb sliding window.

(M) Pearson correlation coefficients between LDB1-AID ChIP-seq replicates

(N) Micro-C contact matrices from merged replicates performed in LDB1-AID cells in untreated and 4 hour auxin-treated conditions. Matrices are shown at three resolutions and window sizes to highlight compartments (left) domains (middle) and loops (right). LDB1 ChIP-seq tracks are shown for untreated and 4 hour auxin-treated conditions.

(O) Schematic of putative loop analysis. APA plots for putative CRE loops that were missed by cooltools loop calling in untreated and 4 hour auxin-treated conditions. 1k resolution.

**Figure S2. LDB1 regulates nascent transcription, related to Figure 2**

(A) TT-seq PCA analysis n=3.

(B) Bar plots showing fold changes for differentially expressed genes identified by TT-seq. Left graph shows the fold change for each gene calculated by RT-qPCR, bottom graph shows the fold change for each gene calculated by TT-seq. For RT-qPCR experiments nascent transcript levels were measured for each gene relative to nascent Gapdh levels. Fold changes relative to the untreated control were calculated for each technical replicate and averaged. Average fold change values are plotted for each biological replicate (n=3). For TT-seq, DESEQ2 normalized counts within gene bodies were measured for each replicate. A fold-change value is plotted for each biological replicate relative to the respective untreated control (n=3).

(C) Pol2 ChIP-seq signal at TSS/TES regions and traveling ratios. Signal at each window and traveling ratios were calculated before/after 4hr of LDB1 depletion.

(D) Boxplots representing the average change in TT-seq signal in gene bodies upon LDB1 depletion. Genes are categorized by the number of LDB1-dependent, putative CRE loop anchors overlapping a 1kb window flanking their TSS. Whiskers represent 10^th^ and 90^th^ percentiles; P-values calculated using a two-sided Mann-Whitney U test.

(E) Fractional stacked bar graph showing the proportion of LDB1-dependent genes with various LDB1 occupancy annotations. LDB1-dependent genes are grouped into 4 mutually exclusive categories based on LDB1 occupancy: 1 – genes with LDB1 binding within 1kb of their TSS (upstream or downstream depending on the direction of transcription), 2 – genes with intronic LDB1 peaks, 3 – genes with LDB1 peaks at exons but no intronic peaks, and 4 – genes with extragenic LDB1 peaks (greater than 1kb away from their TSS, but no peaks in introns or exons).

(F) Heatmaps showing Pearson correlation coefficients among Pol2 ChIP-seq replicates performed in LDB1-AID cells. Heatmaps separately shown for untreated and 4 hour auxin-treated conditions. Pearson correlation coefficients were calculated genome-wide using 10k bins.

**Figure S3. LDB1 mediates small loops identified by RCMC, related to Figure 3**

(A) RCMC (top-right) and Micro-C (bottom-left) Contact matrices (chr8:124,780,000-124,870,000) at ZFPM1 locus. 1k resolution.

(B) Boxplots showing the change in loop strength for loops identified in RCMC using Cooltools. Loops are stratified by LDB1 occupancy: LDB1 unoccupied (left), LDB1 present in one anchor (middle), and LDB1 present in both anchors (right).

(C) Boxplots showing the average change in TT-seq signal in gene bodies upon LDB1 depletion. Genes are categorized by the number of LDB1-dependent loop anchors overlapping a 1kb window flanking their TSS. Loops identified in RCMC using Cooltools. Only genes within captured regions are shown on graph.

(D) RCMC contact matrices at CBFA2T3 locus (500bp resolution) for untreated and 4 hour auxin-treated conditions. Right matrix is zoomed in on CBFA2T3 promoter region. ChIP-seq tracks for LDB1 (red), CTCF (green) and RAD21 (blue) are shown below matrix for untreated and 4 hour auxin-treated conditions.

(E) RCMC contact matrices at BCL2L1 locus (150bp resolution) for untreated and 4 hour auxin-treated conditions. ChIP-seq tracks for LDB1 (red), CTCF (green), and RAD21 (Blue) are shown below matrix for untreated and 4 hour auxin-treated conditions.

**Figure S4. LDB1 occupancy is uncoupled from that of CTCF, YY1 and cohesin, related to Figure 4**

(A) ChIP-seq heatmaps and average profile plots showing CTCF, RAD21 and YY1 ChIP-seq signal in LDB1-AID cells in LDB1 replete and depleted conditions. Signal is only shown at peaks that overlap an LDB1 peak in LDB1 replete conditions for each factor.

(B) Stacked fractional bar plots showing the proportion of weakened RAD21 peaks (weakened upon LDB1 depletion) that overlap LDB1-occupied enhancers or LDB1-unoccupied enhancers (left). Reciprocally, stacked fractional bar plot showing the proportion of LDB1-occupied enhancers that overlap a weakened RAD21 peak (right). Weakened RAD21 peak defined as at least a 50% reduction in RAD21 ChIP-seq signal upon LDB1 depletion.

(C) Numbers of ChIP-seq peak changes for LDB1, RAD21, YY1, and CTCF upon LDB1, SMC3, CTCF and YY1 depletion.

(D) Proportion of weakened CRE loops (identified by Micro-C upon LDB1 depletion) that have a weakened RAD21, YY1 or CTCF peak present in one, or both anchors.

(E) Proportion of strengthened CRE loops with strengthened RAD21, YY1 or CTCF peaks (upon LDB1 depletion) at one or both anchors.

(F) Proportion of weakened CRE loops with strengthened YY1 peaks (upon LDB1 depletion) in one or both anchors.

(G) Pearson correlation coefficients for SMC3-AID and CTCF-AID ChIP-seq replicates

**Figure S5. LDB1 can function in the absence of CTCF and YY1, related to Figure 5**

(A) Focus on dual sensitive CRE loops. (left) pie chart showing number of weakened RAD21 ChIP-seq peaks (upon LDB1 depletion) in anchors of SMC3/LDB1 dual sensitive CRE loops. Overlap of SMC3/LDB1 dual sensitive loops vs CTCF/LDB1 dual sensitive loops. LDB1 ChIP-seq signal at LDB1 peaks in SMC3/LDB1 dual sensitive loops, LDB1 ChIP-seq signal at LDB1 peaks in CTCF/LDB1 dual sensitive loops.

(B) Boxplots showing the change in loop strength for LDB1-dependent CRE loops in response to LDB1 depletion (darker colors) or CTCF depletion (lighter colors). Loops are categorized as LDB1 only loops (red) or dual sensitive loops (green).

(C) Boxplots showing the average H3K27ac ChIP-seq signal at enhancers within LDB1-only loop anchors or dually-sensitive (CTCF and LDB1-dependent) loop anchors. Only mutually exclusive enhancer elements between the two sets are considered.

(D) Boxplots showing the distance to encompassing structural loop anchors for LDB1-only loops and dually sensitive (CTCF and LDB1-dependent) loops. Only loops with an encompassing structural loop are shown.

(E) Boxplots showing the change in loop strength for LDB1-dependent CRE loops in response to LDB1 depletion (darker colors) or YY1 depletion (lighter colors). Loops are categorized as LDB1 only loops (red) or dual sensitive loops (orange).

(F) Boxplots showing the average H3K27ac ChIP-seq signal at enhancers within LDB1-only loop anchors or dually-sensitive (YY1 and LDB1-dependent) loop anchors. Only mutually exclusive enhancer elements between the two sets are considered.

(G) Boxplots showing the distance to encompassing structural loop anchors for LDB1-only loops and dually sensitive (YY1 and LDB1-dependent) loops. Only loops with an encompassing structural loop are shown.

(H) Distribution of loop lengths for LDB1 only and CTCF or YY1/LDB1 dual sensitive CRE loops. Maximum loop length shown 1Mb.

**Figure S6. Mitotic LDB1 ChIP-seq, related to Figure 6**

(A) Representative FACS plots and example gates (black boxes) showing the strategy for isolating mitotic populations. One set of plots representative of three independent biological replicates is shown.

(B) Bar plots showing LDB1 enrichment during each cell cycle stage at a strong LDB1 peak by ChIP-qPCR for biological replicate 1. Enrichment is plotted as a fraction of input material. LDB1 enrichment is compared to an isotype-matched IgG negative control.

(C) ChIP-seq heatmaps for LDB1 at each cell cycle stage for all 3 biological replicates. Heatmaps show LDB1 ChIP-seq signal at all LDB1 peaks identified in asynchronous cells.

(D) Heatmap showing Pearson correlation coefficients between all LDB1 mitotic ChIP-seq samples. Pearson correlation coefficients were calculated using the average RPM signal within peaks identified in asynchronous cells. As expected, a lower concordance amongst replicates is observed for samples with lower signal-to-noise ratios (prometaphase and ana/telophase).

## ACKNOWLEDGEMENTS

We thank Mustafa Mir, Rajan Jain, Douglas Epstein, and members of the Blobel lab for helpful discussions. We also thank the Children’s Hospital of Philadelphia Flow Cytometry Core for assistance with cell sorting. This work was supported by grants T32GM008216 and the Blavatnik Family Fellowship Award to N.G.A.; T32HG000046 and F30DK132824 to J.C.L.; R24DK106766 to R.C.H. and G.A.B.; National Science Foundation of China Grant 321004422 to H.Z.; and R01DK05937, R01DK058044, and U01DK127405 to G.A.B.

## AUTHOR CONTRIBUTIONS

G.A.B. conceived the study. G.A.B. and N.G.A. designed experiments. N.G.A. created the LDB1 auxin-inducible degron cell line used in this study. H.Z. created the SMC3 and CTCF auxin-inducible degron cell lines used in this study. J.C.L. created the YY1 auxin-inducible cell line generated in this study. N.G.A. performed Micro-C, TRI-C, and Pol2 ChIP-seq experiments in the LDB1-AID degron cell line. N.G.A., S.C.M, and A.Q. performed cell cycle LDB1 ChIP-seq experiments. X.W. performed the engineered forced looping experiments in the SMC3-AID cell line. S.W. performed TT-seq experiments. RCMC experiments were designed by A.S.H., V.Y.G., N.G.A., and G.A.B. N.G.A treated and prepped samples for RCMC, V.Y.G performed RCMC protocol. S.C.M. performed ChIP-seq experiments in LDB1-AID, CTCF-AID, SMC3-AID, and YY1-AID cell lines, J.C.L. processed the ChIP-seq data with help from S.C.M. and N.G.A. C.A.K., B.M.G., and R.C.H. contributed to sequencing of ChIP-seq, Micro-C, TT-seq, TRI-C, and RNA-seq. J.C.L. processed Micro-C data. Data analysis was performed by N.G.A. with help from J.C.L. N.G.A. and G.A.B. wrote the manuscript with inputs from all authors.

## STAR METHODS

### RESOURCE AVAILABILITY

#### Lead contact

Further information and requests for resources and reagents should be directed to and will be fulfilled by Gerd A. Blobel.

#### Materials availability

Unique/stable regents or cell lines generated in this study are available upon request to the lead contact.

#### Experimental model and subject details

The G1E-ER4^79^ murine erythroblast cell line was gifted by Dr. Mitchel Weiss. G1E-ER4 cells express GATA1 fused to the ligand binding domain of the estrogen receptor. Addition of 100nM estradiol activates GATA1 and induces erythroid maturation.

### METHODS DETAILS

#### Cell culture and maintenance

The G1E-ER4 cell line and its sublines were cultured in IMDM supplemented with 2% penicillin/streptomycin, 15% fetal bovine serum, Kit ligand, erythropoietin, and monothioglycerol. Cells were maintained at a density less than 1 million cells per 1mL.

#### Generating LDB1-AID cell line

We homozygously inserted minimal-AID (mAID) and mCherry at the endogenous LDB1 locus in G1E-ER4 cells using CRISPR-mediated homology directed repair. We used a donor template designed to insert mAID-mCherry in-frame with the 3’ end of LDB1. The donor template included 900bp of 5’ homology, mAID, mCherry and 939bp of 3’ homology. These sequences were assembled into a vector backbone for cloning purposes using the Takara In-Fusion HD Cloning kit (Takara, 639648). The repair template was then amplified from the cloning vector and purified using QIAquick Gel Extraction Kit (QIAGEN, 28704). Two gRNA sequences each targeting the 3’ end of LDB1 were separately cloned into the px458-GFP plasmid. The purified repair template and the px458-GFP plasmid (containing the LDB1 gRNA and Cas9) were electroporated into G1E-ER4 cells using the Amaxa II electroporator (Lonza) with the Amaxa II Cell Line Nucleofector Kit R (Lonza, VCA-1001). Two separate reactions were performed; one for each gRNA. 6ug of linear repair template and 18ug of px458-GFP plasmid were used in the transfection reactions. After 24 hours, mCherry positive cells were selected by FACS and expanded as single-cell clones. PCR screening was used to identify 2 clones (one from each gRNA reaction) with homozygous insertions of mAID-mCherry. We confirmed editing via Sanger sequencing in each clone. OsTiR-IRES-GFP was expressed in each LDB1-AID cell line with the MigR1 retroviral vector. LDB1-AID cells expressing OsTiR-IRES-GFP were isolated by FACS.

#### Validation of LDB1 depletion upon auxin treatment

LDB1-AID-mCherry G1E-ER4 cells expressing OsTiR-IRES-GFP were treated with 1mM auxin (indole 3-acetic acid sodium salt, Sigma, I5148) for 0, 1, 2, or 4 hours and fixed with 1% formaldehyde. Cells were subjected to flow cytometry to measure mCherry signal. Wildtype G1E-ER4 cells were used as a negative control. To further validate the LDB1-AID response to auxin and compare tagged LDB1 protein levels to untagged LDB1 in the parental line, we performed Western blot analysis for LDB1 (Thermo Fisher Scientific, PA5-56948) in both LDB1-AID clonal lines and parental G1E-ER4 cells in the absence of auxin and in 4 hour auxin treatment conditions. Samples were lysed in complete RIPA lysis buffer and sonicated with the Bioruptor Pico (Diagenode, 3 min: 30sec on, 30sec off, ‘easy’ mode. Protein lysates were run on a 4-12% Bis-Tris gel. GAPDH was used as a loading control (Santa Cruz Biotechnology, sc-32233). RT-qPCR was used to further validate LDB1-AID clonal lines and test their ability to differentiate upon treatment with estradiol. Briefly, RNA from parental G1E-ER4 cells and LDB1-AID clones was isolated using the RNeasy Mini Kit (QIAGEN, 74104). RNA was isolated from cells under the following treatment conditions: untreated, 24 hour treatment with estradiol (Sigma, E2758), simultaneous 24 hour treatment with estradiol and auxin). Genomic DNA was removed from samples using the QiAshredder (QIAGEN, 79656) and on-column digestion with RNAse-free DNAse (provided with RNeasy Mini Kit). cDNA was generated using iSCRIPT Reverse Transcription Supermix (Bio Rad, 1708840). qPCR reaction was performed using SYBR Green PCR Master Mix (Thermo Fisher, 4367660).

#### Micro-C

Micro-C was performed as previously described^75,76^ with minor adjustments. 5 million cells were used as input for each reaction. To increase library diversity, dinucleosomes from 2-3 technical replicates were pooled after gel extraction (prior to library preparation). In brief, cells were crosslinked with 1% formaldehyde for 10 min followed by an additional fixation with 3mM DSG (ProteoChem, c1104-1gm) for 40 min. Fixed cells were permeabilized with Micro-C Buffer 1 at a concentration of 1 million cells/100uL (50 mM NaCl, 10 mM Tric-HCl (pH 7.5), 5 mM MgCl2, 1 mM CaCl2, 0.2% NP-40, 1 X Protease Inhibitor Cocktail tablet (Millipore Sigma, 11836170001)) for 20 min on ice. Chromatin from permeabilized nuclei was digested with 10 U MNase (Worthington Biochemical, LS004798) for 10 min at 37C with 850rpm rotation. Digested fragments were de-phosphorylated with 5 U r-SAP (New England Biolabs, M0371S) for 45 min at 37C in de-phosphorylation buffer (50mM NaCl, 10mM Tris-HCl, 10mM MgCl2, 100 ug/mL BSA). De-phosphorylated fragments were subjected to end-chewing using 20 U T4 PNK (New England Biolabs, M0201S) and 40 U large Klenow Fragment (New England Biolabs, M0210S) for 15 min at 37C in the following buffer: 50mM NaCl, 10 mM Tris-HCl, 10 mM MgCl2, 100 ug/mL BSA, 2 mM ATP, and 3 mM DTT. Biotin incorporation was achieved by adding biotin-dATP (Jena Bioscience, NU-835-BIO14-S), biotin-dCTP (Jena Bioscience, NU-809-BIOX-S), dTTP, and dGTP and incubating at 25C for 45 min. Finally, fragmented and labeled DNA ends were ligated using 5,000 U of T4 DNA ligase (New England Biolabs, M0202S) and incubating at room temperature for 180 min with rotation. Unligated ends were removed by exonuclease III for 10 min at 37C. After reverse-crosslinking, DNA was purified using PCI and ethanol precipitation and size selected for dinucleosmal fragments by gel extraction. Informative fragments were immobilized on MyONE Strptavidin C1 Dynabeads (Thermo Fisher, 65001). Sequencing libraries were prepared using NEBNext Ultra II DNA Library Prep Kit with NEBNext unique dual index primer pairs and amplified with KAPA HiFi Hot Start Mix (Roche, 08202940001). 9 biological replicates per treatment condition were sequenced (2×50bp) on the Illumina Nextseq platform.

#### RNA-seq

RNA was isolated from parental G1E-ER4 cells and LDB1-AID clones using the RNeasy Mini Kit (QIAGEN, 74104) according to manufacturer’s specifications. Genomic DNA was removed from samples using the QiAshredder (QIAGEN, 79656) and on-column digestion with RNAse-free DNAse (provided with RNeasy Mini Kit) according to manufacturer’s specifications. Sequencing libraries were constructed from 500 ng of DNase-treated, total RNA using the TruSeq Stranded mRNA kit (Illumina cat# 20020594) for polyA+ selection, cDNA synthesis and library preparation according to manufacturer’s specifications. Briefly, first strand cDNA was synthesized from polyA+ selected RNA using reverse transcriptase and random primers, followed by second strand synthesis, end repair, 3’ adenylation, and adaptor ligation. Completed libraries were amplified by PCR for 11 cycles. The quality and size (mean 318 bp) of each library was evaluated using the Agilent Bioanalyzer 2100 using the DNA 7500 kit (cat# 5067-1504), followed by quantitation using real-time PCR using the KAPA Library Quant Kit for Illumina (KAPA Biosystems catalog no. KK4835). Libraries were then pooled and sequenced in paired-end mode using a P2 flow cell on the NextSeq 2000 to generate 2 x 76 bp reads using Illumina-supplied kits as appropriate. FASTQ were demultiplexed using Illumina’s DRAGEN Bio IT Platform v3.7.4 and sequence reads were processed using the ENCODE3 long RNA-Seq pipeline (https://www.encodeproject.org/pipelines/ENCPL002LPE/). In brief, reads were mapped to the mouse genome (mm9 assembly, GENCODE vM1 genes) using STAR, followed by RSEM for gene quantifications.

#### ChIP-seq

Chromatin immunoprecipitation (ChIP) was performed using the following antibodies: Pol2 (Cell Signaling, D8L4Y, 10uL/IP), LDB1 (Santa Cruz, sc-365074, 10ug/IP), CTCF (Millipore, 07-729, 10ug/IP), RAD21 (Abcam, ab992, 10ug/IP), YY1 (Active motif, 61779, 10ug/IP). In brief, cells were lysed in 1 ml ice-cold cell lysis buffer (10 mM Tris pH 8, 10 mM NaCl, 0.2% Igepal) supplemented with protease inhibitors and PMSF) for 20 min. Nuclei were pelleted and lysed using 1 mL Nuclear Lysis Buffer (50 mM Tris pH 8, 10 mM EDTA, 1% SDS) supplemented with PI and PMSF for 20 min on ice. Samples were sonicated with the Bioruptor Pico (Diagenode, 5 min: 30sec on, 30sec off, ‘easy’ mode). Nuclear extracts were precleared with 50uL protein A/G agarose beads (Thermo Fisher, 15918014 and 15920010) and 50 ug isotype-matched IgG for at least 2 hours. 200uL of chromatin was taken as input. Chromatin was incubated with 35 uL A/G beads that were pre-bound with antibody (10ug/IP) and incubated at 4C overnight. Beads were washed one time with IP wash buffer I (20mM Tris pH 8, 2 mM EDTA, 50 mM NaCl, 1% Triton X-100, 0.1% SDS), twice with high-salt buffer (20 mM Tris pH 8, 2 mM EDTA, 500 mM NaCl, 1% Triton X-100, 0.01% SDS), once with IP was buffer 2 (10 mM Tris pH 8, 1 mM EDTA, 0.25 M LiCl, 1% Igepal, 1% NA-deoxycholate), and twice with TE buffer (10 mM Tris pH 8, 1 mM EDTA). All washes were performed with ice-cold buffers on ice. Beads were then moved to room temperature and eluted in 200 ul using elution buffer (100 mM NaHCO3, 1% SDS). 2 uL RNAseA (10mg/ml) and 12 ul 5M NaCl were added to input and IP samples and incubated at 37C for 30 min. 3 uL of 20mg/ml proteinase K was added and samples were reverse crosslinked at 65C overnight. 10 uL of 3 M sodium acetate was added to all samples and DNA was purified using QiAquick PCR purification kit (QIAGEN, 28104). ChIP-seq libraries were prepared using NEBNext Ultra II DNA Library Prep Kit with NEBNext unique dual index primer pairs. Libraries were sequenced (2×50bp) on an Illumina NextSeq 500 platform. Pol2 ChIP-seq libraries were sequenced (1×75bp).

#### TT-seq

TT-seq was performed as previously described^82,108^. Exponentially growing cells were labeled with 500 μM 4-thiouridine (4SU) (MedChemExpress), for 5 minutes. Cells were processed with 2 mL TRIzol Reagent (Invitrogen) (per 10 million G1E-ER4 cells) and total RNA was extracted following manufacturers instructions. 500 ng of 4SU-labeled Drosophila Schneider 2 (S2) cells total RNA was used as spike in and was mixed with 100 μg of collected 4SU-labeled G1E-ER4 total RNA. Mixed RNA was fragmentated using a final concentration of 0.2 M NaOH for 18 minutes and neutralized with 0.5 M Tris-HCl (pH 6.8). RNA was purified by isopropanol precipitation. Labelled RNA was biotinylated in 300 μL of biotinylation mix (fragmented total RNA, 10 mM HEPES pH 7.5, 1 mM EDTA, 0.167 mg/mL MTSEA-biotin (Biotium)) for 1 hour at room temperature and purified with phenol/chloroform/isoamyl alcohol (25:24:1) extraction. Denaturation of biotinylated RNA was carried out at 65 °C for 10 minutes, followed by rapid cooling on ice for 5 minutes. The denatured biotinylated RNA was bound to Dynabeads MyOne Streptavidin C1 (Invitrogen) at room temperature for 30 minutes, eluted with 100 mM DTT and purified by isopropanol precipitation. RNA quality was determined using Agilent TapeStation RNA ScreenTape (Agilent). Strand-specific sequencing libraries were generated using the Illumina Stranded Total RNA Prep (Illumina) and IDT for Illumina RNA UD Indexes Set A, Ligation (Illumina). Library size was determined using Agilent TapeStation High Sensitivity DNA ScreenTape (Agilent). Libraries were pooled and sequenced on the Illumina NextSeq 500 platform.

#### RCMC

RCMC was performed as previously described^37^. The RCMC protocol merges Micro-C (described above) with region capture via tiling of biotinylated probes. Target loci were selected based on the presence of LDB1-dependent genes (identified via TT-seq/Pol2 ChIP) and genomic features of interest. For example, Myc was selected as it is an LDB1-dependent gene within a gene-poor TAD. An added advantage of this locus is that LDB1-occupied enhancers within the Myc TAD are relatively widely spaced and thus some LDB1-dependent CRE loops involving Myc are detectable by Micro-C, allowing us to validate RCMC findings. Conversely, we also selected LDB1-dependent genes within gene-dense regions (eg. Zfpm1 and Cbfa2t3). Micro-C lacks the resolution to detect loops between closely-spaced LDB1 peaks at the Zfpm1 and Cbfa2t3 loci. We selected roughly 1-Mb-sized regions that included loci of interest. 80-mer biotinylated probes were designed to tile end-to-end with no overlap across the capture regions through Twist Bioscience. Probes in high-repeat regions were removed from the probe tiling. Probes were synthesized and purchased as Custom Target Enrichment Panels from Twist Bioscience. Capture was performed using Twist Bioscience’s Standard Hybridization target Enrichment Protocol. Libraries were dried and mixed with Hybridization Mix (Twist Bioscience, 104178), Custom Target enrichment Panels and Universal Blockers (Twist Bioscience, 100578), along with Mouse Cot-1 DNA (Thermo Fisher, 18440016). Hybridization was carried out overnight. Pull down was performed with streptavidin beads (Twist Bioscience, 100983) which were subsequently washed (Twist Bioscience, 104178). Target-enriched libraries were PCR amplified using Equinox Library Amplification Mix (Twist Bioscience, 104178). Libraries were purified (Twist Bioscience, 100983) and sequenced (2×50) on an Illumina NovaSeq 6000 system. RCMC data in this paper was generated from two biological replicates. A list of the 5 captured loci (mm9 coordinates) are provided in Table S3.

#### Tri-C

TRI-C was performed as previously described^78,90^ with minor modifications. 15 Million cells were used for each replicate. A total of 4 biological replicates were performed for each treatment condition. Cells were fixed with 2% formaldehyde for 10 min at room temperature. Formaldehyde crosslinking was quenched with 0.125M glycine. Cells were permeabilized for 20 min on ice in 5 mL of cold cell lysis buffer (10 mM Tric-HCl pH 8, 10 mM NaCl, 0.2% Igepal, 1x EDTA-free cOmplete Protease Inhibitor cocktail). Permeabilized cells were resuspended in 1 mL of cold PBS and flash frozen with liquid nitrogen. Fixed cells were thawed on ice, spun for 15 min at 500 X g, 4C and resuspended in 650 uL 1XNlaIII restriction buffer. Cells were split into 3 aliquots (200uL each) and the following were added sequentially to each aliquot: 404 uL nuclease-free water, 60uL 10x NlaIII restriction buffer, 10uL 20% SDS. The remaining 50 uL of fixed nuclei was used as nondigested control. All tubes were shaken at 37C at 500 rpm (intermittent: 30s on/30 s off) for 1 HR. 66 uL of 20% Triton X-100 was added to each digestion reaction and incubated for another 1 HR. 300 U NlaIII (New England Biolabs, R0125L) was added until the end of the day, an additional 300 U were added overnight (37C at 500rpm intermittent shaking). An additional 250U of NlaIII was added to each digestion and incubated at 37C 500rpm intermittent shaking for an additional 6 HRs. 100 uL was taking from each digestion reaction and saved as nonligated control. NlaIII was heat inactivated at 65C for 20 min and immediately cooled on ice. 642 uL ligation solution (0.4 U/uL T4 DNA ligase in 2.1X T4 DNA ligase buffer) was added to each reaction and incubated at 16C, 500rpm (intermittent 30s on/30s off shaking) for ∼22HRs. Ligation reactions were centrifuged at 500 X g for 15 min and nuclei were resuspended in 300uL TE buffer. 5 uL of 600U/mL proteinase K was added to each reaction and incubated at 65C overnight. 5 uL of 15 U/mL RNAse A was added to each ligation reaction and incubated at 37C for 30 min. DNA was extracted using standard phenol-chloroform-isoamyl alcohol and ethanol precipitation. Ligation efficiency was estimated by running controls and 5-10uL of 3C library on a 1% agarose gel. 3C library was quantified using Qubit dsDNA BR assay (Thermo Fisher, Q32850). 6 ug of 3C library was sonicated and split into 2 NEBNext reactions for library preparation. Samples were sonicated to 400-500bp fragments using the Bioruptor Pico (Diagenode, 2min: 30sec on, 30sec off, ‘ultralow’). Sonicated 3C libraries were purified with 0.7X AMPure XP beads (Beckman Coulter, A63880). Sonicated material was split into 2 aliquots and 2 NEBNext reactions were performed per sample for library prep. End Prep, adaptor ligation and USER enzyme steps were performed as per manufacturors instructions. DNA was amplified using Herculase II DNA polymerase (Agilent, 600675) and mixed dual index primers. Amplified libraries were purified with 1.8X ampure XP beads. Oligonucleotide capture was performed using KAPA HyperCapture Reagents (Roch, 9075810001). Capture steps were multiplexed such that 1 oligonucleotide capture was performed in a pooled fashion for multiple uniquely indexed libraries in a single tube. Uniquely indexed libraries were pooled at 1:1 mass ratio for each capture reaction (1-2ug was used for each library). 5ug/library of mouse C0t DNA (Thermo Fisher, 18440016) was added to the DNA pool. Complex was concentrated using vacuum centrifuge eat 50C until sample was completely dry. 6.7uL per library of universal enhancing oligonucleotides was added to resuspend desiccated DNA. 14uL per library of 2X Hybridization buffer and 6uL of Hybridization Component H was added to the mixture and incubated at room temperature for 2 min. 4.5 uL per library of biotinylated capture oligonucelotide targeting the Myc promoter region was added and sample was transferred to a thermocycler and incubated at 95C for 5 min and then 47 C for 72 HRs. 50 uL per library of Dynabeads M-270 streptavidin beads (Thermo Fisher, 65305) were used to enrich for captured DNA. Beads were washed with 1 X Bead Wash buffer and placed on a magnetic stand to remove supernatant. Beads were resuspended with the hybridization reaction and bead/library complex was incubated at 47C for 45 min with 600 rpm shaking. 50 uL per library of 1X Wash buffer I was added to the beads and bound DNA and placed on a magnetic stand. Supernatant was discarded. Beads were subsequently washed with 100 uL per library of Stringent Wash Buffer (pre-heated to 47C) twice (incubated at 47 for 5 min after each wash). Beads were washed with 100 uL per library room temperature Wash buffer I, then subsequently with 100uL per library of wash buffer II (room temperature) and finally with 100 uL of room temperature Wash buffer III. Beads were resuspended in PCR-grade water and captured DNA was amplified (off the beads) using KAPA HiFi Hot Start Ready mix with capture primers and supernatant was purified with 1.8X ampure XP beads. A second capture step was performed to further enrich for our region of interest similar to the first. For the second capture, volumes for hybridization reaction and bead washing were added for a single library and hybridization reaction occurred for ∼22Hrs. Finally, DNA libraries were sequenced on the illumina (NEXTseq platform, 2×150bp).

#### Isolating mitotic populations via FACS

We utilized a G1E-ER4 subline expressing mCherry-MD for mitotic LDB1 ChIP-seq experiments. mCherry is fused to the mitotic degradation domain of cyclin B and thus specific cell populations can be isolated based on mCherry signal and DNA content during the mitosis-G1 transition. The sorting method and cell line were described previously^36^. Briefly, cells were treated with 200ng/mL of nocodazole for 8.5 hours. Cells were either collected at 8.5 hours of nocodazole treatment to enrich for prometaphase cells or were pelleted, washed with warm, nocodazole-free media and released for the following timepoints to enrich for different populations during the mitosis-G1 transition: 25min (ana/telophase), 1 hour (early G1), 2 hours (mid G1) or 4 hours (late G1). After harvesting each cell population, cells were cross-linked with 1% formaldehyde. Cross-linking was quenched with 1M glycine, and cells were permeabilized with 0.1% TritonX-100. All samples were stained with 0.5ul/10 million cells anti-pMPM2 antibody (Millipore, 05-368) for 50 min at RT. Secondary antibody staining was performed with APC-conjugated F9ab’)2-Goat anti-Mouse (Thermo Fisher Scientific, 17-4010-82) for 30 min at RT. Finally, cells were resuspended in FACS buffer supplemented with 25ng/mL DAPI and kept on ice. Cells were subject to flow sorting on the MoFlo Astrios EQ sorter (Beckman Coulter). Prometaphase samples were sorted based on positive mCherry-MD, positive pMPM2 and 4N DAPI signal. Ana/telophase samples were sorted based on 4N DAPI signal and reduced mCherry-MD signal. Early G1, mid G1, and late G1 samples were sorted on 2N DAPI signal and negative mCherry-MD signal. Sorted cells were aliquoted and flash frozen. We performed 3 biological replicates of ChIP-seq for LDB1 at each of the cell cycle stages. Representative FACS plots and example gating strategies for each cell cycle population are shown in Figure S6A.

#### Micro-C data processing and visualization

We used the distiller pipeline (v3.3) to generate contact maps using fastq files as input. PCR duplicates were removed from each replicate and balanced contact maps were generated for each treatment condition from merged biological replicates. Iterative correction and eigenvector decomposition (ICE) balancing was used to normalize contact maps using default settings: a given bin was excluded if its sum was >5 median absolute deviations below the median bin, the first two diagonals were ignored for balancing, columns and rows were normalized so that they summed to 1. We used coolbox^109^ (v0.3.8) to visualize contact maps and aligned ChIP-seq tracks. To generate pileup plots (APA plots) of Micro-C contacts, we used cooltools (v0.5.3) to average contact frequencies across loops.

#### Micro-C compartment analysis

We used cooltools (v0.5.3) to compute cis eigenvector values from 100kb binned matrices from untreated and auxin-treated conditions. We generated saddle plots which reflected all AA, BB, AB, BA interactions.

#### Micro-C domain analysis

We followed a similar approach outlined in (Zhang et al., 2021) to call domains. Briefly, we identified domains using the rGMAP^86^ software using 10kb-binned contact matrices. We then generated a final merged and filtered domain list using the following strategy: we merged domain calls from untreated and auxin-treated samples, removed duplicate domains, merged domains that had start and end coordinates within 80kb of each other, removed domains that were smaller than 100kb and larger than 2mb. We defined boundaries as 120kb windows flanking the start/end positions of each domain. We calculated insulation scores at boundaries using cooltools with a 120kb sliding window. Finally, we analyzed insulation scores at domain boundaries by calculating the minimum insulation score at all boundaries for untreated and auxin-treated samples. Our final merged and filtered domain list from rGMAP was used to identify CRE loops within/across TADs in Figure 2 and to assess gene expression changes based on LDB1-occupied enhancer density within TADs. We recapitulated these results using an independent TAD caller: HiTAD from TADLib^87,88^. The results using HiTAD-identified domains show the same trends we observed using rGMAP-identified domains.

#### Loop calling and quantification

To identify and quantify loops, we used the approach outlined in Lam et al. (manuscript under review). Cooltools.dots was used to identify loops using merged contact maps for each treatment condition. We first identified loops on 2kb, 5kb, and 10kb resolution contact maps separately for each treatment condition using the following parameters: max_loci_separation=2_000_000, clustering_radius=20_000, lambda_bin_fdr=0.05 and n_lambda_bins=50. Default settings were used to define dots (pixels) enriched relative to local neighborhoods: donut, vertical, horizontal, and lowleft. However, we utilized rounded donut and lowleft neighborhoods to more easily identify loops close to the diagonal. We created a master loop list by merging untreated and auxin-treated loops from each resolution. Redundant loops were merged (redundancy defined as being the same pixel or adjacent pixels). Then, we merged consensus lists from each resolution (2kb, 5kb, and 10kb), retaining the smallest resolution coordinates in instances where a loop was called at multiple resolutions. Loop strength was quantified by calculating the observed/locally-adjusted expected value. The locally-adjusted expected value was calculated by multiplying the expected value at the loop’s peak pixel by the sum of the observed contacts in the rounded donut region divided by the sum of the expected contacts in the rounded donut region. Loop strength was calculated for each loop using the resolution at which the loop was identified. Loops with strengths of 0, NA, or infinite were removed to filter out loops in sparse regions. This resulted in a final list of 20,926 chromatin loops. We then calculated the log2FC for loop strength such that negative values reflected loops that were weakened upon auxin treatment and positive values reflected loops that were strengthened upon auxin treatment. We used a log2FC cutoff of −/+ 0.5 to define weakened/strengthened loops. A similar strategy as described above was used to call and quantify loops for RCMC, except we used the following resolutions to call loops for RCMC data: 500bp, 1kb, 2kb, and 5kb, and we used the following clustering radii cutoffs respectively: 1_000, 2_000, 4_000, and 10_000. RCMC allows for the identification of loops at higher resolutions and can more accurately distinguish between adjacent loops compared to Micro-C.

#### Characterizing loops based on ChIP-seq peaks and CRE annotations

We used previously-annotated sets of putative active enhancers and promoters (Zhang et al., 2019) based on merged H3K27ac ChIP-seq peaks in uninduced G1E-ER4 cells. Putative active promoters were defined as H3K27ac peaks within 1kb of a TSS, putative active enhancers were defined as H3K27ac peaks greater than 1kb away from a TSS. To characterize loops, we created fixed loop anchors of 10kb (by adding/subtracting 5kb from the original anchor center). Then, we intersected putative active enhancers/promoters, LDB1 ChIP-seq peaks, and CTCF/RAD21 peaks (defined as RAD21 peaks with at least one bp overlap with a CTCF peak) with loop anchors using bedtools^110^ intersect with the -c flag. We then characterized loops based on the presence/absence of CREs and CTCF/RAD21 peaks into the following categories: CRE loops – loops with an enhancer or promoter at both anchors, structural loops – loops with CTCF/RAD21 in both anchors and not CRE at both anchors. Note, structural loops can contain a CRE in one anchor but not both (see table S4 for all annotated Micro-C loops). We further stratified CRE loops into 4 additional subcategories: enhancer/enhancer loops – enhancers at both anchors but no promoters at either anchor, promoter/promoter loops – promoters in both anchors but not enhancers at either anchor, enhancer/promoter loops -enhancer at one anchor and promoter at the opposite anchor (these loops cannot have enhancer and promoter in the same anchor), and mixed loops – have enhancer and promoter in the same anchor and thus cannot be classified into the other subcategories. When we use the term “LDB1-dependent CRE loop” these are loops with LDB1 present in at least one anchor, an enhancer or promoter at both anchors and are weakened (log2FC < −0.5) upon LDB1 depletion.

#### Integrating looping changes from multiple degron cell lines

To determine whether LDB1-dependent loops were also dependent on CTCF, cohesin, or YY1, we calculated the loop strength of all loops identified from our Micro-C data sets using published 10kb-binned CTCF-AID HiC contact maps from G1E-ER4 cells isolated in mid G1 phase (Zhang et. al., 2021), 10kb-binned HiC contact maps from SMC3-AID asynchronous G1E-ER4 cells (Zhao et. al., under review), and 10kb-binned Micro-C contact maps from YY1-AID asynchronous G1E-ER4 cells (Lam et. al., under review). We quantified loop strength for untreated and auxin-treated samples for all loops identified using the LDB1-AID degron cells and removed loops with 0, NA, or infinite loop strength values to filter out loops in sparse regions. We then calculated log2FC values to reflect changes in looping with respect to CTCF, SMC3, or YY1 depletion. To determine whether LDB1-dependent CRE loops were also dependent upon CTCF, cohesin, or YY1 we identified loops with an LDB1 peak in at least one anchor, had an enhancer or promoter at both anchors and were weakened in the LDB1-AID system (log2FC < −0.5). We then determined the number of these loops that were either sensitive (log2FC < −0.5) to CTCF/cohesin/YY1 degradation or resistant (log2FC > −0.5) to CTCF/cohesin/YY1 degradation.

#### Integrating transcription with looping

To integrate transcription with looping, we combined our chromatin looping data with our TT-seq data. We defined 1kb windows centered on the TSSs of genes. We intersected the anchors of loops with these TSS windows and categorized genes by the number of loop anchors that overlapped. We split genes into 3 categories based on their loop interactions: genes that did not interact with any loop, genes that interact with 1 loop, and genes that interact with 2 or more loops. We did so separately for 4 mutually exclusive loop types: LDB1 dependent CRE loops (CRE loops with an LDB1 ChIP-seq peak in at least one anchor and weakened upon LDB1 depletion), LDB1 independent CRE loops (CRE loops with no LDB1 ChIP-seq peak in either anchor and unchanged upon LDB1 depletion), strengthened CRE loops (CRE loops strengthened in the absence of LDB1), and strengthened nonCRE loops (loops with CRE at one or no anchors and strengthened upon LDB1 depletion). We then analyzed the log2FC values for genes in each category. Gene Log2FC values were calculated using DESeq2 and represent the average change of TT-seq read counts within gene bodies from 3 biological replicates. Before integrating with looping, genes were removed that had a Padj value set to NA. DESeq2 assigns Padj NA values to genes with low read counts or contain a sample with an extreme outlier based on Cook’s distance. We used default DESeq2 settings to identify outliers and define low read counts. In addition to measuring the average fold change for genes connected to loops, we also measured their baseline expression levels. To do so, we calculated the average CPM-normalized, strand-specific TT-seq signal across each gene body using bwtool^111^ summary. This gives the average signal normalized for gene length.

#### ChIP-seq data processing and analysis

ChIP-seq was performed for 2-3 biological replicates for each cell line, IP, and treatment condition. Input material corresponding to each cell line and treatment condition were also sequenced. Reads were aligned to the mm9 reference genome using Bowtie2^112^ (2.4.5). Duplicate reads were filtered out using SAMtools^113^ (1.3.1) with MAPQ<20. We generated bigwig files for each replicate using deeptools^114^ (v3.5.1) bamCoverage. After confirming concordance amongst replicates, we generated summary bigwig files for each IP/treatment condition by merging replicate bigwig files. We did so in one of two ways: 1-for ChIP-seq experiments using the LDB1-AID and CTCF-AID cell lines, we had 2 replicates for each sample allowing us to use the deeptools bamCompare function to average the signal from each replicate and create summary, BPM-normalized bigwigs (--binSize 20, --normalizeUsing BPM, --operation mean), 2 -for ChIP-seq experiments using the SMC3-AID cell line, cell cycle ChIP-seq experiments and Pol2 ChIP experiments, we had 3 replicates for each sample, we merged BAM files using Samtools merge and generated BPM (or CPM for Pol2 ChIP)-normalized bigwig files from the merged BAM files using deeptools bamCoverage (--binsize 20, --normalizeUsing BPM or CPM). Deeptools computeMatrix and plotHeatmap were used to generate heatmaps and profiles of ChIP-seq signals. Macs2^115^ (v2.2.9.1) was used to call narrow peaks for each replicate using the paired-end setting and a matched input bam file as a control. The peak-calling threshold was set to p = 1e-5. Peaks were combined from each replicate, centered and set to standardized 400bp regions. For cell cycle LDB1 ChIP-seq experiments, we generated heatmaps and profiles for ChIP-seq signals at LDB1 peaks identified in asynchronous cells. For LDB1-AID, CTCF-AID, YY1-AID, and SMC3-AID data, heatmaps and profiles were generated for ChIP-seq signal at peaks identified in the untreated condition. To test the concordance amongst ChIP-seq replicates, we used deeptools multiBigwigSummary (in bins mode) and plotCorrelation to calculate pearson correlations between all samples using 10kb genomic bins.

#### TRIC data processing and analysis

The Capcruncher pipeline is an all-in-one data processing pipeline for TRI-C and Capture-c experiments. Capcruncher was used to process TRI-C data using the -TRI option. The capcurncher pipeline takes raw fastq files as input and using the TRI-C option, will filter uniquely mapped reads for those containing a capture site (in our case, to ensure all filtered contacts have one fragment overlapping the MYC capture probe) and at least 2 ligation junctions (to ensure all filtered reads represent multi-way contacts). 1 kb binned contact matrices from 4 biological replicates were merged for each treatment condition using cooler^116^ merge with the -join and -Header flags to generate merged.cool files for untreated and auxin-treated samples. 5Kb binned contact matrices were generated using cooler zoomify on the merged cool files and on the individual replicate files. Raw contacts were corrected for the number of NLAIII restriction fragments in each bin-bin pair. Each matrix was then scaled to a total of 1 million contacts so that direct comparisons can be made between conditions. Normalized contact matrices on the merged files were visualized using coolbox. Normalized contacts for each individual replicate were used to quantify the number of multi-way contacts involving LDB1 ChIP-seq peaks.

To quantify multiway contacts involving LDB1, we filtered replicate cool files (binned at 5kb resolution) to retain contacts where both interacting bins overlapped an LDB1 chip-seq peak and summed all LDB1-LDB1 multiway contacts for each replicate and each treatment condition. We removed multi-way contacts where the interaction occurred within the same bin as to correctly identify multiway contacts driven by distinct LDB1-bound sites. Importantly, we do not detect an LDB1 chip-seq peak at the Myc promoter region, thus all multiway contacts involving distinct LDB1-bound bins represent multiway contacts between distinct LDB1-occupied sites and the Myc promoter. We performed the same analysis except filtering for contacts that did not contain LDB1 peaks in either interacting bin as a control.

#### TT-seq data processing and analysis

TT-seq paired-end reads were trimmed using Trim Galore (v0.6.10) and mapped to the mouse mm9 reference genome using STAR v2.7.10b. Reads with MAPQ smaller than 7 were filtered out and duplicate reads were marked using SAMtools v1.14 or Picard v3.0.0. Strand-specific TT-seq reads in gene bodies were quantified using deepTools v3.5.1. DESEQ2^117^ was used to perform differential expression analysis.

