## Supplementary figures and images for "LDB1 establishes multi-enhancer networks to regulate gene expression"

### Supplemental Figures

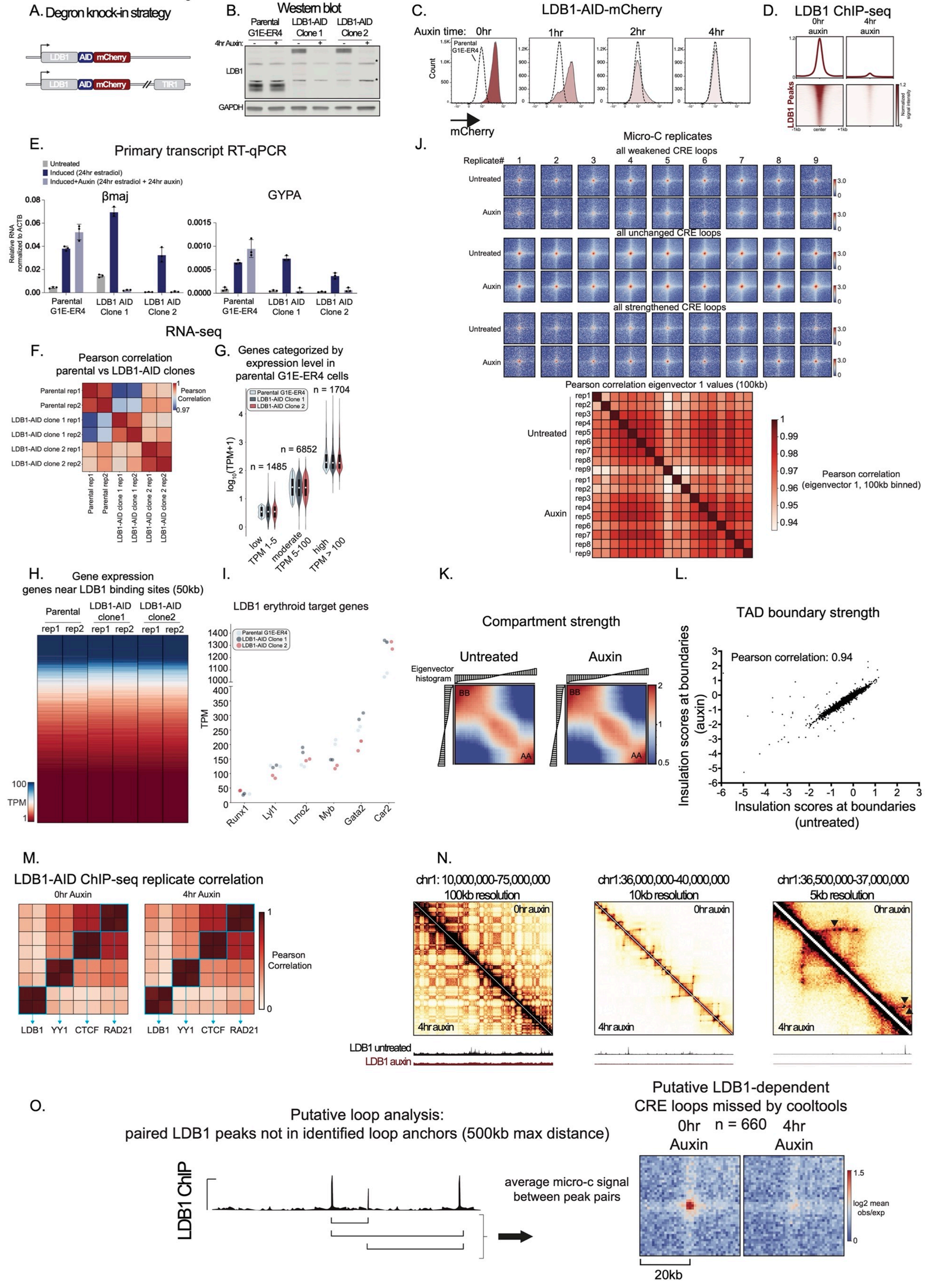

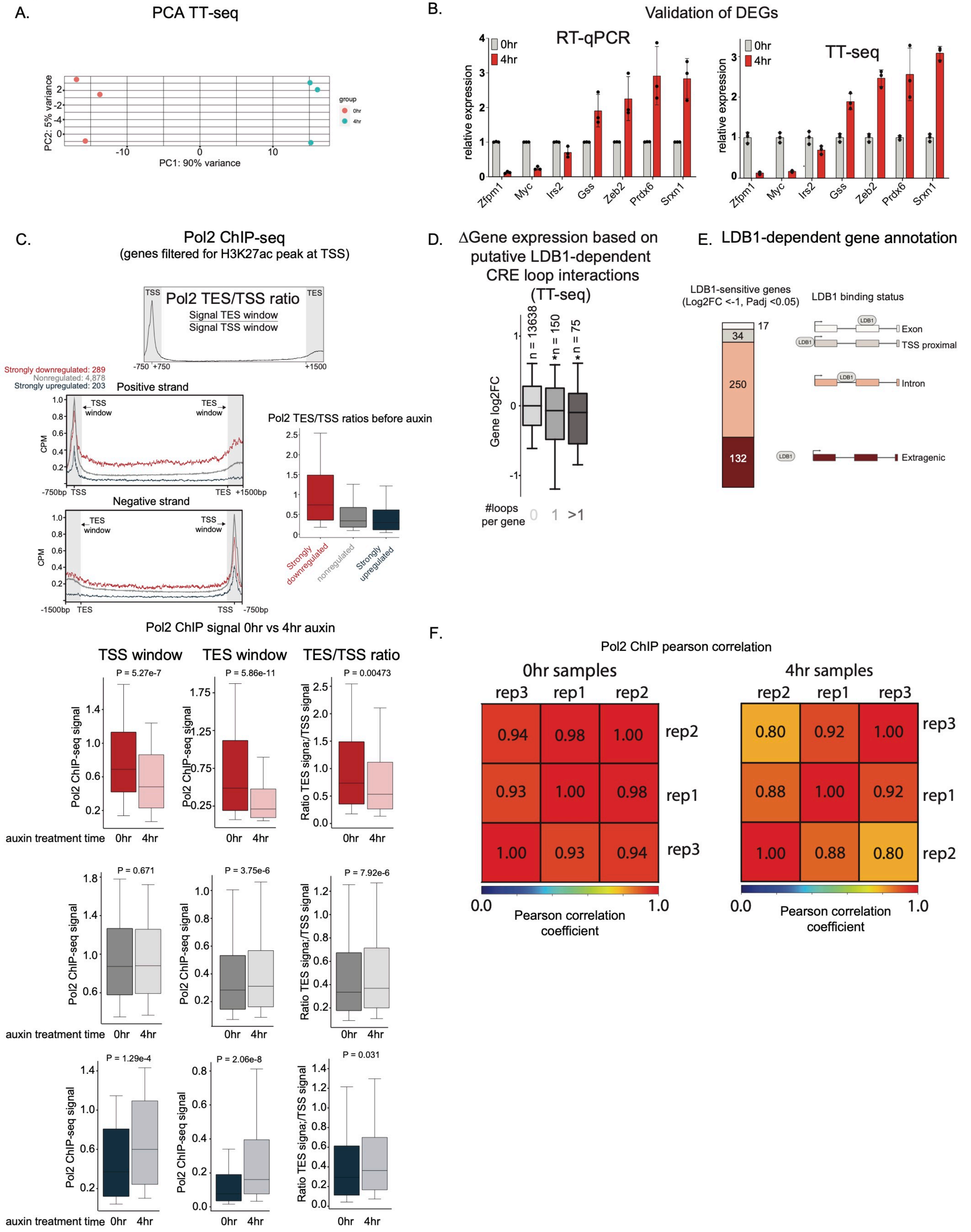

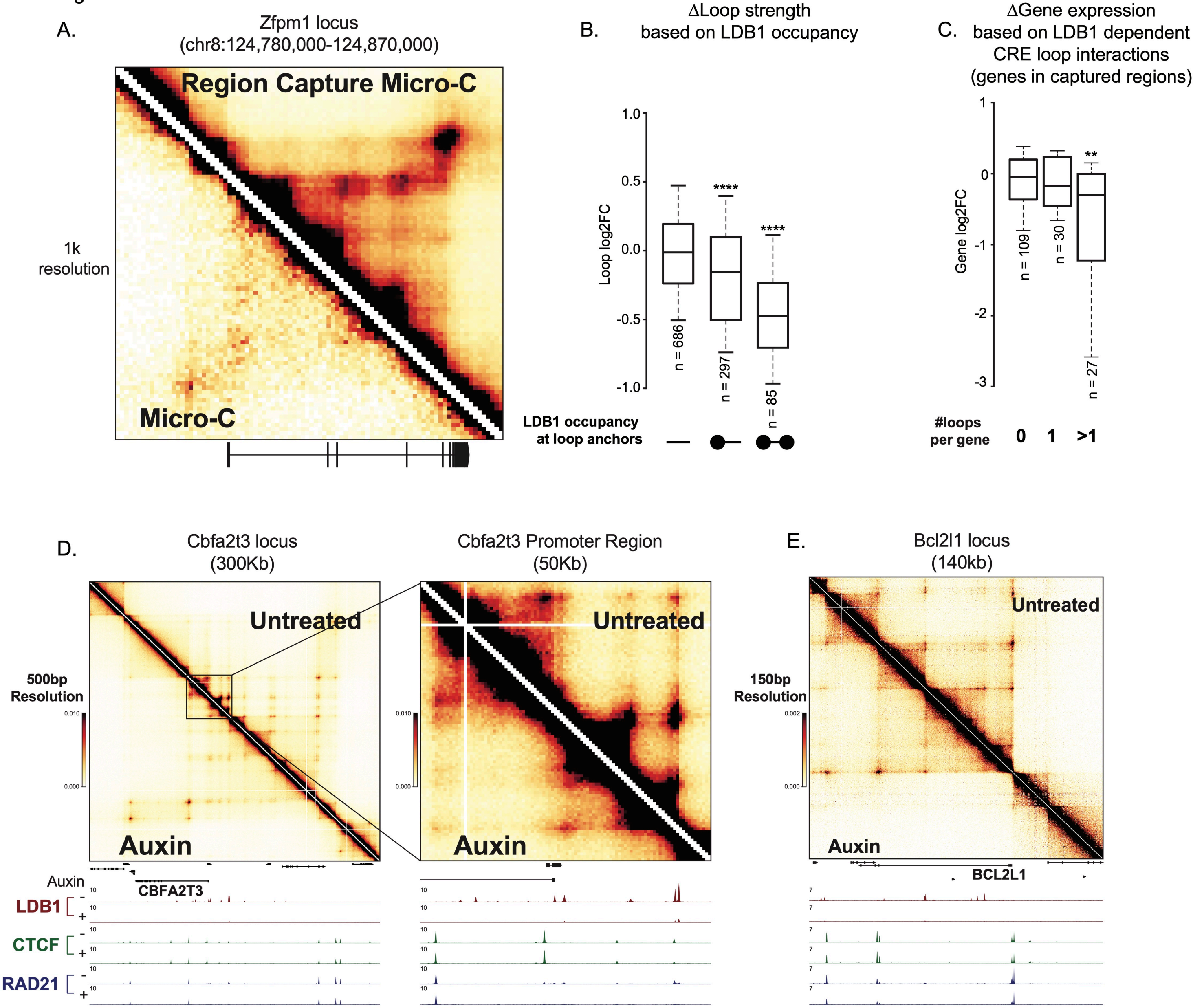

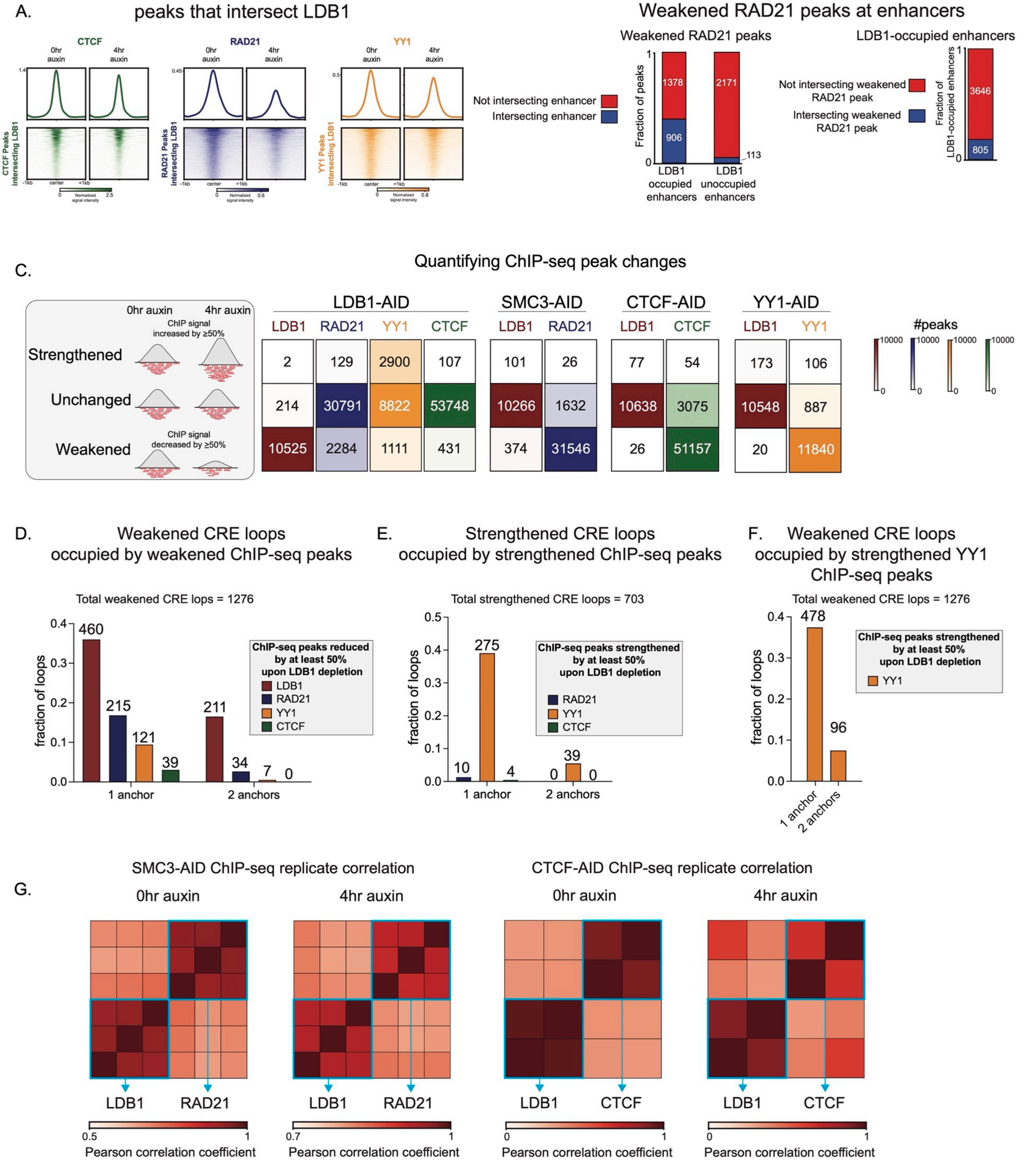

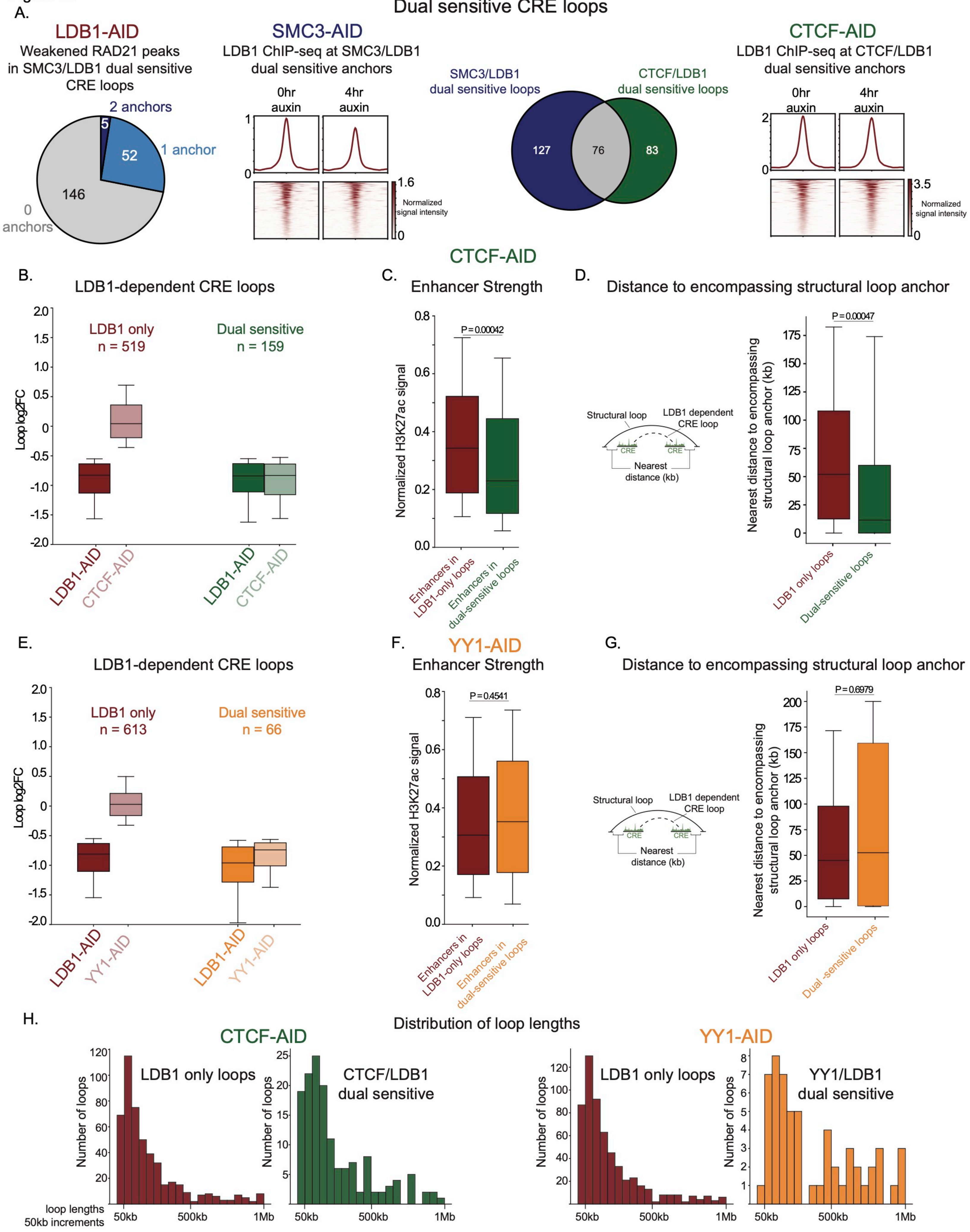

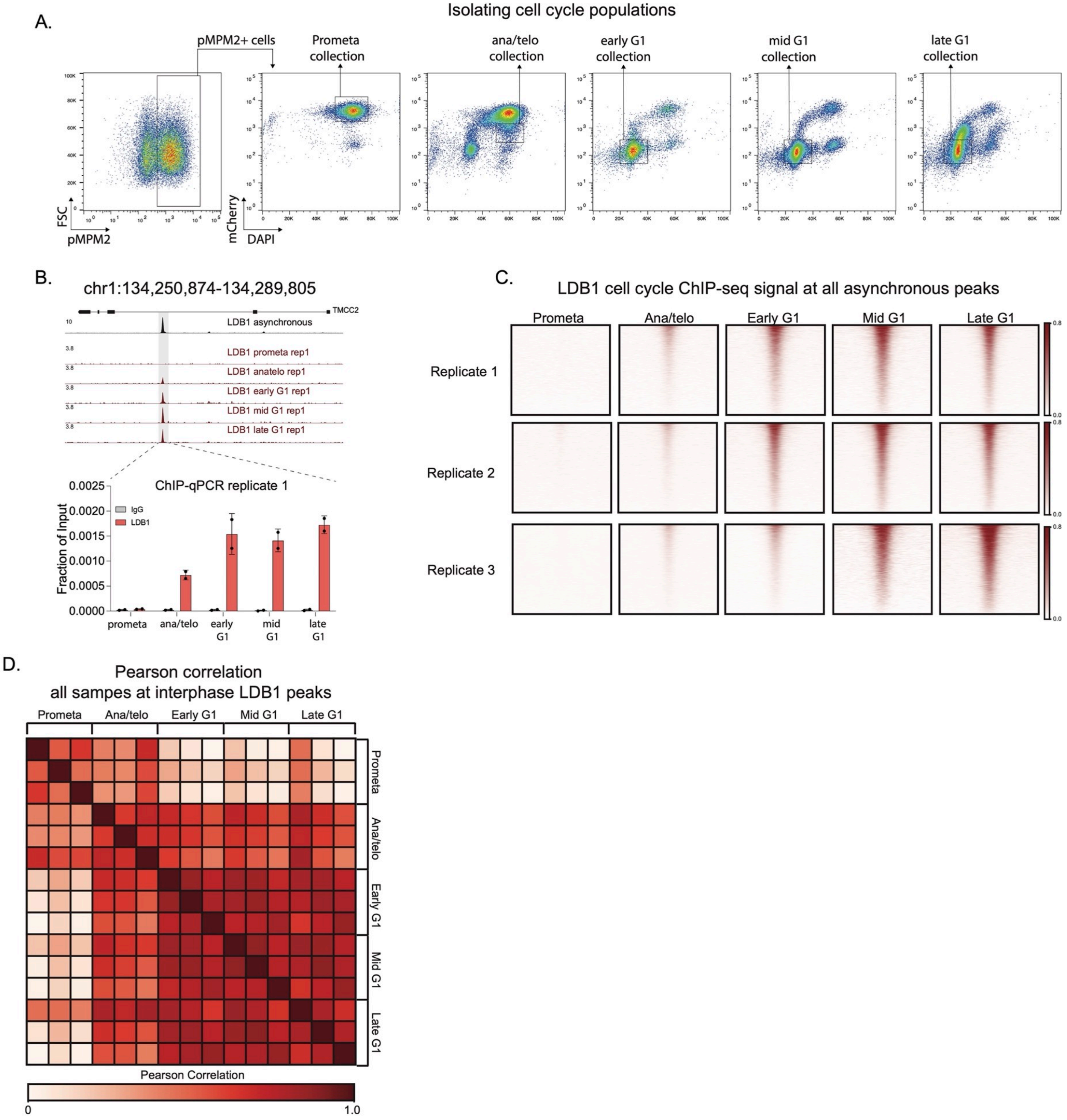
